# Intrinsic p53 Activation Restricts Gammaherpesvirus-Driven Germinal Center B Cell Expansion during Latency Establishment

**DOI:** 10.1101/2023.10.31.563188

**Authors:** Shana M. Owens, Jeffrey M. Sifford, Gang Li, Steven J. Murdock, Eduardo Salinas, Mark Manzano, Debopam Ghosh, Jason S. Stumhofer, J. Craig Forrest

**Affiliations:** Dept. of Microbiology and Immunology and Center for Microbial Pathogenesis and Host Inflammatory Responses; Winthrop P. Rockefeller Cancer Institute, University of Arkansas for Medical Sciences, Little Rock, AR 72205

**Author notes:** J. Craig Forrest, Dept. of Microbiology and Immunology, University of Arkansas for Medical Sciences, 4301 W. Markham, Slot 511, Little Rock, AR 72205. These authors contributed equally to this work (Shana M. Owens, Jeffrey M. Sifford).

## Abstract

Gammaherpesviruses (GHV) are DNA tumor viruses that establish lifelong latent infections in lymphocytes. For viruses such as Epstein-Barr virus (EBV) and murine gammaherpesvirus 68 (MHV68), this is accomplished through a viral gene-expression program that promotes cellular proliferation and differentiation, especially of germinal center (GC) B cells. Intrinsic host mechanisms that control virus-driven cellular expansion are incompletely defined. Using a small-animal model of GHV pathogenesis, we demonstrate *in vivo* that tumor suppressor p53 is activated specifically in B cells that are latently infected by MHV68. In the absence of p53, the early expansion of MHV68 latency was greatly increased, especially in GC B cells, a cell-type whose proliferation was conversely restricted by p53. We identify the B cell-specific latency gene M2, a viral promoter of GC B cell differentiation, as a viral protein sufficient to elicit a p53-dependent anti-proliferative response caused by Src-family kinase activation. We further demonstrate that EBV-encoded latent membrane protein 1 (LMP1) similarly triggers a p53 response in primary B cells. Our data highlight a model in which GHV latency gene-expression programs that promote B cell proliferation and differentiation to facilitate viral colonization of the host trigger aberrant cellular proliferation that is controlled by p53.

**IMPORTANCE:** Gammaherpesviruses cause lifelong infections of their hosts, commonly referred to as latency, that can lead to cancer. Latency establishment benefits from the functions of viral proteins that augment and amplify B cell activation, proliferation, and differentiation signals. In uninfected cells, off-schedule cellular differentiation would typically trigger anti-proliferative responses by effector proteins known as tumor suppressors. However, tumor suppressor responses to gammaherpesvirus manipulation of cellular processes remain understudied, especially those that occur during latency establishment in a living organism. Here we identify p53, a tumor suppressor commonly mutated in cancer, as a host factor that limits virus-driven B cell proliferation and differentiation, and thus, viral colonization of a host. We demonstrate that p53 activation occurs in response to viral latency proteins that induce B cell activation. This work informs a gap in our understanding of intrinsic cellular defense mechanisms that restrict lifelong GHV infection.

## INTRODUCTION

Gammaherpesviruses (GHVs) are DNA tumor viruses that include the human pathogens Epstein-Barr virus (EBV) and Kaposi sarcoma-associated herpesvirus (KSHV) and the rodent pathogen murine gammaherpesvirus 68 (MHV68), among others. Following a period of productive viral replication in various host tissues, GHVs establish lifelong chronic infections^1,2^. This phase of the infectious cycle, referred to as latency, is maintained in B cells. The lifelong infections caused by GHVs place the host at risk for numerous cancers, including Burkitt lymphoma, Hodgkin lymphoma, primary effusion lymphoma, multicentric Castleman disease, and many others^3,4^. Moreover, GHV-related malignancies represent serious complications for AIDS patients and transplant recipients^5^. However, the precise mechanisms by which GHVs promote oncogenesis are incompletely defined, as are the host pathways that prevent cellular transformation.

To establish and maintain latency, GHVs stimulate and usurp normal B cell differentiation processes^6^. This is accomplished through the actions of viral noncoding RNAs and oncoproteins, which, upon infection of naïve B cells, lead to an activated phenotype reminiscent of a germinal center (GC) reaction^7,8^. GC B cells are highly proliferative, with an estimated doubling time of every 6 hours^9^. GC B cells also undergo mutagenic processes known as somatic hypermutation and class-switch recombination, which change the antigenic specificity and effector functions of the B cell receptor and secreted antibodies^10^. The GC reaction ultimately results in the emergence of long-lived memory B cells or antibody-secreting plasma cells that respectively enable recall responses upon reinfection and facilitate elimination of pathogens^11^. It is hypothesized that stimulating GC reactions enables GHV-infected B cells to increase in number dramatically as the cells rapidly proliferate while also allowing a mechanism for establishment of long-term latency in the circulating memory B cell pool^12^. This model of GHV latency establishment is largely based on EBV infection and immortalization of primary B cells in culture, but it is also supported by kinetic evaluations of cell-types that harbor MHV68 *in vivo* and through analyses of mutant viruses following experimental infection of mice [reviewed in ^13,14^].

For instance, the MHV68 homolog of the latency-associated nuclear antigen (mLANA) encoded by *ORF73* is essential for latency establishment after intranasal inoculation of mice^15–17^. In addition to mediating maintenance of the viral episome during latent cell division, mLANA also promotes c-Myc stabilization, which likely influences latent cell survival^18^. The *ORF72*-encoded viral cyclin D ortholog (v-Cyclin) is a *bona fide* oncogene^19^, and it is important for viral replication and reactivation from latency^20,21^. A viral Bcl-2 homolog (v-Bcl-2) encoded by the *M11* open-reading frame, prevents activation-induced cell death to support viral latency^22–24^. The *M2*-derived gene product is a scaffold protein that in-and-of-itself promotes B cell differentiation by activating Src family kinases, phospholipase C-gamma (PLCγ), and downstream transcription via nuclear factor of activated T cells (NFAT) and interferon regulatory factor 4 (IRF4)^25–28^. Interestingly, M2’s role in facilitating MHV68 latency establishment appears to function almost exclusively in the GC B cell compartment^29,30^. EBV likewise uses a coordinated gene expression program that facilitates latency establishment and promotes cellular proliferation and differentiation^1^. Of note, latent membrane protein 1 (LMP1) mimics the activating and pro-survival signaling functions of CD40^31–33^, a B cell surface receptor that interacts with CD40L on T follicular helper cells to facilitate GC reactions^10^. M2 and LMP1 are hypothesized to be functional homologs despite no discernable sequence homology^14^.

While many of the viral oncogenes that promote cellular proliferation, survival, and differentiation have been studied extensively, less is known regarding the potential for cellular tumor suppressor pathways to be activated in response to manipulation of host-cell physiology by GHV proteins during latency establishment. It is noteworthy that EBV, KSHV, and MHV68 each encode viral proteins capable of inducing cell-cycle entry and transformation^19,32,34–36^, yet none of these viruses efficiently transforms infected cells. This suggests that the viral latency-associated gene expression programs induce host cell responses that limit the potential for cellular transformation during infection. For example, although most if not all cells are infected and begin to proliferate after EBV infection of primary B cells in culture, only a small percentage of infected cells immortalize to become lymphoblastoid cell lines (LCLs)^37–40^. This is because EBV-induced cellular proliferation triggers the activation of ataxia telangiectasia mutated (ATM) and checkpoint kinase 2 (Chk2), DNA damage response (DDR) kinases that inhibit cellular proliferation and subsequent immortalization^37^. KSHV also triggers ATM activation and proliferative arrest upon infection of primary endothelial cells^41^. Like EBV, MHV68 can immortalize primary cells in culture^42^. Although host restriction factors are not yet known, the process is similarly inefficient, requiring weeks of culture prior to the outgrowth of transformed cells. While DDR pathways appear to restrict GHV-mediated cellular transformation in culture, tumor suppressor responses that restrict GHV infection and immortalization of cells *in vivo* during natural infection of a host are poorly defined and may differ from responses occurring in cell culture. For instance, a role for ATM in restricting MHV68 infection *in vivo* was tested; however, ATM deficiency did not promote enhanced latent MHV68 infection or cancer^43^. Rather, ATM facilitates B cell latency^44,45^, suggesting that ATM does not impose a critical barrier to latent MHV68 infection or related oncogenesis *in vivo*.

Another candidate host protein for intrinsic latency restriction is tumor suppressor p53. p53 is considered a master regulator of genetic integrity, a postulate emphasized by the presence of inactivating p53 mutations in approximately half of all human cancers^46^. Though constitutively expressed, p53 under normal physiologic cellular conditions is constantly targeted for proteolytic degradation by proteins such as ubiquitin ligase mouse double-minute 2 (MDM2)^47^. In response to potentially genotoxic cellular stresses, including DNA damage, oncogene expression, and viral infection, p53 becomes stabilized and activated through a variety of post-translational modifications^47^. Active p53 regulates transcription of numerous cellular genes involved in cell-cycle arrest, DNA repair, and apoptosis^48^. Interestingly, p53 is stabilized and p53-responsive transcripts are induced during B cell immortalization by EBV^49^, and the KSHV-encoded cyclin D ortholog is sufficient in-and-of-itself to induce p53^34,41,50^. Exogenous induction of p53 by MDM2 inhibition reduces LCL formation by EBV^51^, and conversely p53 inhibition facilitates endothelial cell proliferation following KSHV infection^41^. Although both EBV and KSHV encode latency proteins demonstrated to inhibit p53 in various biochemical evaluations^52–54^, established cell lines in which these viral inhibitors of p53 are highly expressed remain responsive to p53 agonists^49,55^, suggesting that these proteins inefficiently suppress p53 functions in latently infected cells. Moreover, it is somewhat counterintuitive that a virus that establishes a life-long chronic infection would universally inhibit a tumor suppressor as critically important as p53. However, p53 mutations do occur frequently in endemic Burkitt lymphoma (eBL), a cancer characterized by stable EBV infection and the presence of chromosomal translocations, which suggests that p53 functions limit GHV-related cancers *in vivo*^2^. Whether p53 is activated to limit GHV infection and tumorigenic potential has not been directly tested using *in vivo* models of pathogenesis.

Here we describe experiments that test the hypothesis that p53 is activated in response to GHV manipulation of B cell physiology during latency establishment. Using MHV68 infection of WT and p53-deficient mice, we define the functions of p53 in controlling viral latency establishment and maintenance *in vivo*. We employed a targeted sufficiency screen using primary B cells to identify specific viral gene products from MHV68 and EBV that are latency-related p53 agonists. Finally, we performed a pharmacologic inhibitor analysis that highlights key signaling events that promote p53 activation. Through this work, we demonstrate that virus-mediated manipulation of B cell activation and differentiation is countered by an intrinsic host tumor suppressor response to limit early proliferative events during latency establishment by a DNA tumor virus.

## RESULTS

### MHV68 induces p53 during latency establishment

Tumor suppressor p53 is activated in response to aberrant cellular proliferation, differentiation, and genotoxic stress, especially when induced by cellular and viral oncogenes ^34,56,57^. In order to establish latency, GHVs encode proteins that promote cellular proliferation and differentiation^3,4,58^, which could hypothetically be recognized by the p53 pathway as potentially oncogenic events. While it is known that p53 is induced by GHV infection of many cell types in culture^41,59^, its activity is putatively inhibited by certain latency gene products^56,57^. However, whether p53 is activated and functional during GHV latency establishment in a living host is not known.

To begin to define the relationship between p53 and GHV latency, we infected mice with an MHV68 recombinant virus that encodes YFP to enable direct identification of infected cells by flow cytometry^60^ and evaluated p53 induction in spleens of infected mice on day 16 after infection. At this time-point acute viral replication has waned, latent viral burdens in the spleen are at their peak, and the majority of infected B cells are proliferating and exhibit a germinal center phenotype^60–62^. In this analysis, p53 levels were significantly higher specifically in the MHV68-infected cells (YFP^+^) than in mock-infected animals or the uninfected cells (YFP^-^) in spleens of infected mice (**Fig. 1a,b, and S1**). Quantitative RT-PCR analyses of bulk B cells detected increased transcription during infection of p53-responsive genes *Cdkn1a* and *Mdm2* (**Fig. 1c**), suggesting that p53 activity was induced during MHV68 latency establishment.

**Figure 1:**
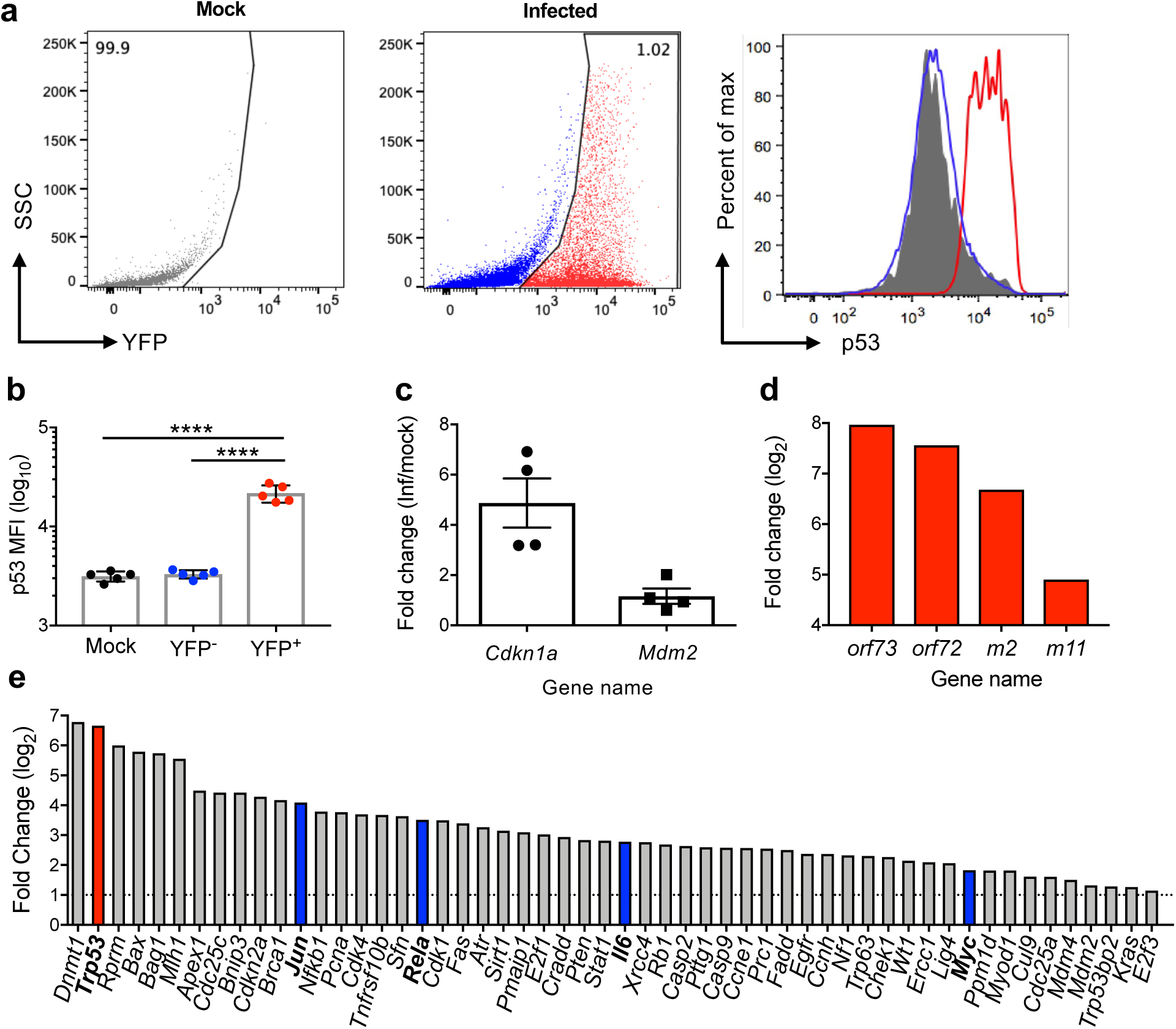
p53 is induced and active in latently infected splenocytes. **a,** Flow cytometry analysis of p53 induction in infected (YFP+, red) and uninfected (YFP-, blue) spleen cells on day 16 after mock-infection (gray) or infection of C57BL/6 mice with 10^4^ PFU of H2B-YFP MHV68. **b,** Quantification of data in **a** represented as mean ± SEM. Mann-Whitney unpaired *t* test, **** p<0.0001. **c,** Quantitative RT-PCR analysis of unsorted splenocytes to detect representative p53-responsive transcripts *cdkn1a* and *mdm2* normalized to *β-actin* on day 16 post-infection. Infection-associated changes were determined by comparison to uninfected splenocytes using the ΔΔ*C_T_* method. **d, e,** Quantitative RT-PCR analyses comparing MHV68 infected to uninfected splenocytes after sorting on day 16 post-infection. The virus-dependent genetic marking strategy to identify infected cells and confirmation of infection are shown in **Supplemental** Figure 1. Viral latency transcript abundance is shown in **d**. Profiler p53 pathway array data are shown in **e**. Relative transcript abundance was determined by comparing virus-positive to virus-negative cells after sorting using the ΔΔ*C_T_* method. The *trp53* transcript is highlighted in red. Transcripts previously evaluated in MHV68 infection are denoted with blue bars.

To more specifically evaluate p53-related transcription within latently infected cells, we infected inducible fluorescent reporter mice^63^ with a recombinant MHV68 virus that expresses Cre recombinase to induce fluorescent protein expression^64,65^ as a means to mark infected cells. After sorting fluorescent cells and confirming infection by limiting-dilution PCR (LD-PCR)^66^ and p53 induction by flow cytometry (**Fig. S1**), we performed p53 pathway RT-PCR array analyses comparing infected cells, which were approximately 99% CD19^+^/B220^+^ B cells (**Fig. S1**), to uninfected B cells from infected animals. Further confirming that the cells were indeed infected, viral latency transcripts were highly enriched in the fluorescent cell population, as were numerous p53-associated transcripts (**Fig. 1d,e**). Of note, *Trp53* transcription was ca. 100-fold higher in infected cells compared to uninfected cells, as were other transcripts with previously demonstrated roles in MHV68 latency, such as *Myc*, *Il6*, and *Nfkb1*^18,67,68^. Together these data support the conclusion that p53 is stabilized and transcriptionally active *in vivo* during MHV68 latency establishment. The observation that p53 stabilization occurred almost exclusively in infected cells using two independent marking strategies demonstrates that p53 activation is not a general consequence of viral infection as might be expected of a typical antiviral response, but rather represents a cell-intrinsic reaction to MHV68 infection.

### MHV68 latency establishment is enhanced in p53-deficient mice

Given that p53 induction was specific to infected cells, we hypothesized that p53 functions in an intrinsic cellular response that controls MHV68 latency establishment. If correct, we predicted that the absence of p53 would correlate with enhanced MHV68 latency establishment. To test this hypothesis, we evaluated MHV68 infection of both p53^+/+^ and p53^-/-^ mice in side-by-side comparisons. Remarkably, on day 16 post-infection the frequency of splenocytes harboring MHV68 genomes was 14-fold higher in p53^-/-^ animals than WT littermates, with approximately 1 in 20 p53^-/-^ cells infected vs. 1 in ca. 280 WT cells (**Fig. 2a,d**). LD-PCR findings were supported by flow cytometry analyses in which nearly 10% of cells in the spleens of p53^-/-^ animals were YFP^+^ (**Fig. 2b**). Moreover, immunohistochemical analyses to detect infected cells also demonstrated an increase relative to WT animals in the number of infected cells present within B cell follicles of p53-null mice (**Fig. 2e**).

**Figure 2:**
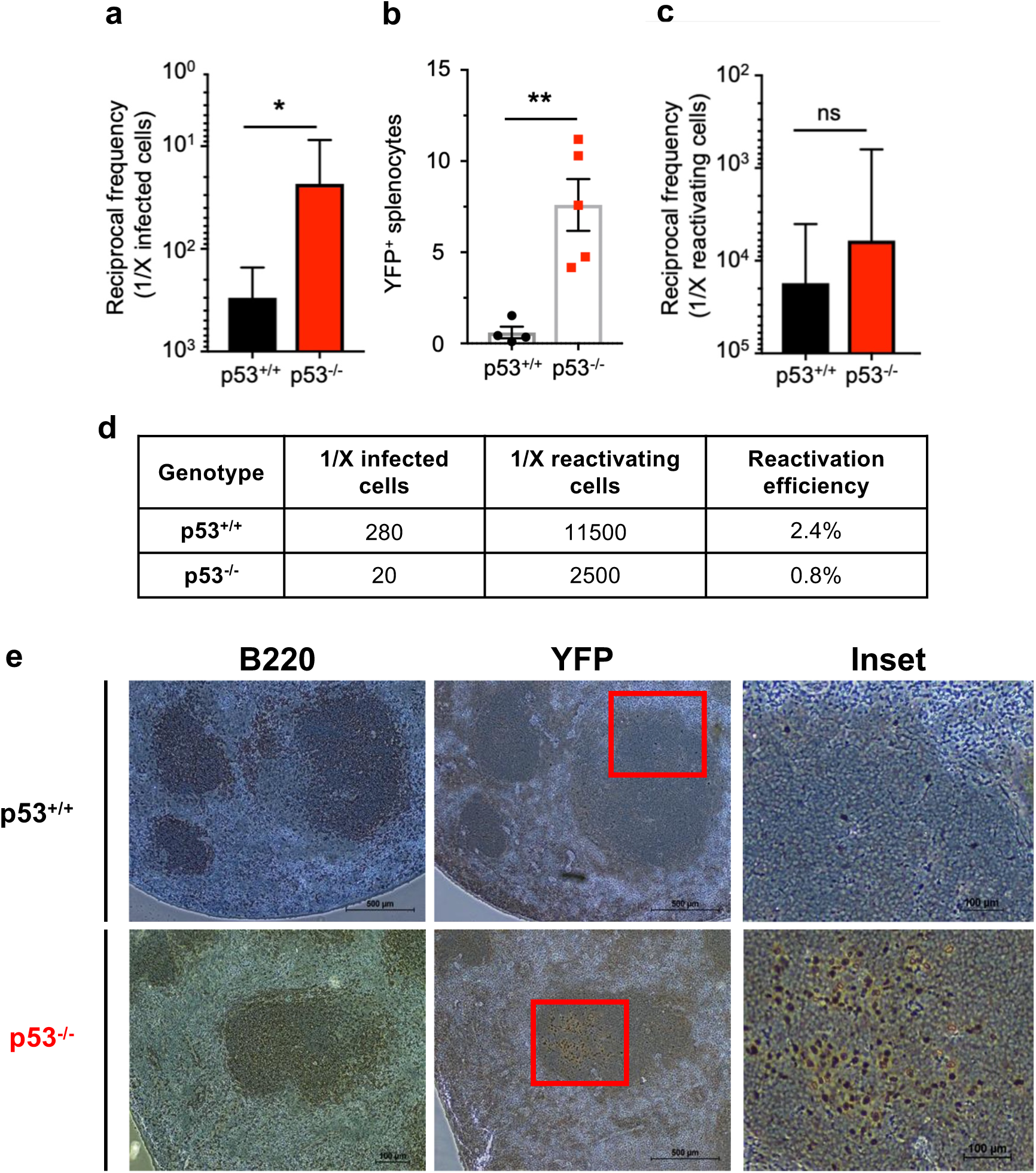
MHV68 latency establishment is enhanced in p53-deficient mice. **a,** Quantification of latently infected splenocyte frequencies by limiting-dilution PCR on day 16 post-infection. p53^-/-^ or p53^+/+^ mice were infected intranasally with 10^4^ PFU of H2B-YFP MHV68. Spearman correlation, * p<0.5 **b,** Flow cytometry analyses to determine the percentage of infected splenocytes (YFP+) for each mouse genotype. Mann-Whitney unpaired *t* test, ** p<0.005. **c,** Quantification of latent MHV68 reactivation efficiency from infected p53^-/-^ or p53^+/+^ splenocytes on day 16 post-infection. Reactivation was detected by scoring cytopathic effect in a limiting-dilution *ex vivo* culture. Data represent means +/- SEM for 2-3 independent infections with a minimum of 3 mice per group. No significant difference by Spearman correlation, ns **d,** Frequencies of infection and reactivation from panels **a,c. e,** Immunohistochemical visualization of B cells (B220+) and MHV68 infected cells (YFP+) in spleen sections from p53^-/-^ or p53^+/+^ mice on day 16 after infection. The inset shows YFP+ cells within a splenic follicle. Increased B220 staining in p53^-/-^ specimens was consistent in multiple sections. Representative images are shown. Scale bars indicate 500 μm or 100 μm for the inset.

Previous studies suggest that p53 plays roles in innate and adaptive immune responses to certain infections^69–71^, so we reasoned mice lacking p53 might be less able to control MHV68 infection. In evaluating acute viral replication, we found no difference in MHV68 infection of WT and p53 knockout mouse lungs on day 7 post-infection (**Fig. S2**), the typical peak of acute viral replication in this tissue^72^. This result suggests that p53 is not critical for controlling the acute phase of MHV68 infection in mice. While the frequency of latently infected splenocytes with the capacity to reactivate upon *ex vivo* culture was enhanced modestly compared to WT mice (**Fig. 2c**), the increase correlated with the higher frequency of cells harboring MHV68 in p53^-/-^ mice (**Fig. 2d**), and we did not detect evidence of ongoing persistent viral replication^66,73^. The absence of increased acute replication, enhanced reactivation, or persistent replication, outcomes largely controlled by innate immune responses^74–78^, suggests that p53 is not directly involved in these host defense mechanisms. Finally, MHV68-specific adaptive immune responses also developed normally in p53^-/-^ mice, as virus-specific IgG in serum, T cell activation, and induction of MHV68 antigen-specific CD8^+^ T cells were comparable in both WT and p53-null animals infected with MHV68 (**Fig. S3**). Together, these data demonstrate that the absence of p53 correlates with enhanced MHV68 latency establishment and support the hypothesis that p53 mediates intrinsic cellular resistance to chronic MHV68 infection.

### p53 controls cellular proliferation during MHV68 latency establishment

As mentioned above, MHV68 and other GHVs encode latency-associated gene products that promote the expansion of B cells that serve as reservoirs for latent infection. For MHV68 this results in a syndrome akin to infectious mononucleosis caused by EBV^79^. In addition to enhanced latent MHV68 infection in p53-deficient animals, it was readily apparent that there was also a significantly greater infection-related increase in the number of cells per spleen compared to WT mice (**Fig. 3a**). This observation suggested that p53 acts to control cellular expansion induced by the virus. In support of this hypothesis, quantification of specific lymphocyte populations by flow cytometry revealed enhanced expansion of B cells in p53^-/-^ mice (**Fig. 3b**), while T cell and natural killer (NK) cell numbers were similar between WT and knockout animals (**Fig. S3**). For particular B cell subsets, the number of GC B cells was significantly increased (**Fig. 3c**), but plasmablasts were only modestly affected by the presence or absence of p53 (**Fig. 3d**). Reflecting MHV68’s latent tropism for B cells and usage of the GC B cell compartment during latency establishment, the number of infected B cells was significantly increased in p53 knockout mice, with the MHV68 infected subset of GC B cells exhibiting a ca. 10-fold relative increase (**Fig. 3e-h**). Infection of plasmablasts also was significantly enhanced, which may result from differentiation of infected GC B cells. In contrast to infection of p53^-/-^ mice with WT virus, infection with a latency-deficient mutant virus lacking the MHV68 homolog of the latency-associated nuclear antigen (mLANA-null) did not cause increased expansion of splenocyte populations (**Fig. S4**). Since this mutant virus can still undergo acute replication and stimulate adaptive immunity^15,16^, these data suggest that the enhanced splenocyte expansion that occurs in p53^-/-^ mice requires the presence of latent MHV68 and is not simply a consequence of the general immune response to viral infection.

**Figure 3:**
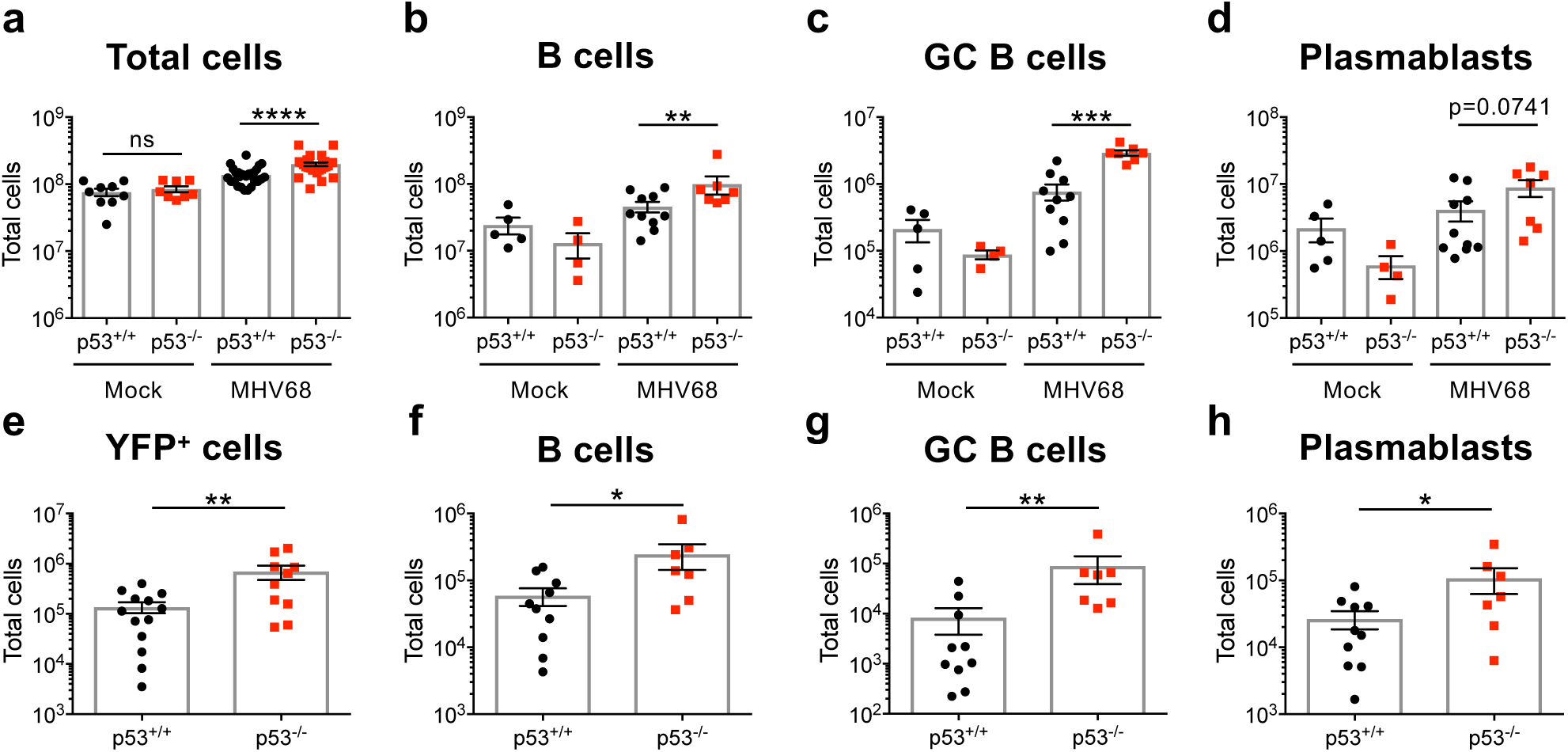
B cell expansion and infection is increased in p53-deficient mice. **a-d,** Quantification of total splenocytes, B cells, GC B cells, and plasmablasts 16 days after mock infection or infection of p53^+/+^ or p53^-/-^ mice with 10^4^ PFU of H2B-YFP MHV68. Flow cytometry was performed to identify specific B cell populations. B cells were gated as CD19^+^/B220^+^ in **b**, GC cells as GL7^+^/CD38^lo^ sub-population of CD19^+^/B220^+^ B cells in **c**, and plasmablasts as CD138^+^/B220^lo^ in **d**. Gating strategies are shown in **Supplemental** Figure 6. Other lymphocyte populations were also quantified and were equivalent in p53^-/-^ and p53^+/+^ spleens (**Supplemental** Figure 3b). **e-h,** Quantification of infected (YFP+) total splenocytes, B cells, GC B cells, and plasmablasts as identified in **a-d** on day 16 after infection. Data represent means +/- SEM for 2-3 independent infections with a minimum of 3 mice per group. Mann-Whitney unpaired *t* test, * p<0.05, ** p<0.005, *** p<0.0005, **** p<0.0001.

Given that GHVs drive B cell proliferation in order to establish and maintain latency^12,58^ and p53 restricts cellular proliferation or promotes apoptosis downstream of oncogene expression^56,80–82^, we hypothesized that p53 functions to limit MHV68-driven cellular proliferation and/or induce death in infected cells. Using *in vivo* 5-ethynyl-2’-deoxyuridine (EdU)-incorporation to label cells actively replicating their DNA, we detected similar basal levels of EdU incorporation in both p53^-/-^ and WT animals that were mock-infected. Infection with MHV68 led to an increase in the number of EdU^+^ GC B cells irrespective of genotype; however, spleens of p53-deficient mice contained twice as many EdU^+^ GC B cells than WT mice (**Fig. 4a**). Cell death, quantified using annexin V and propidium iodide staining, was only modestly reduced in the absence of p53 (**Fig. 4b**). These data suggest that a mechanism by which p53 limits MHV68 latency establishment is through restriction of cellular proliferation, an interpretation supported by the enhanced expansion and infection of the highly proliferative GC B cell compartment in p53^-/-^ mice.

**Figure 4:**
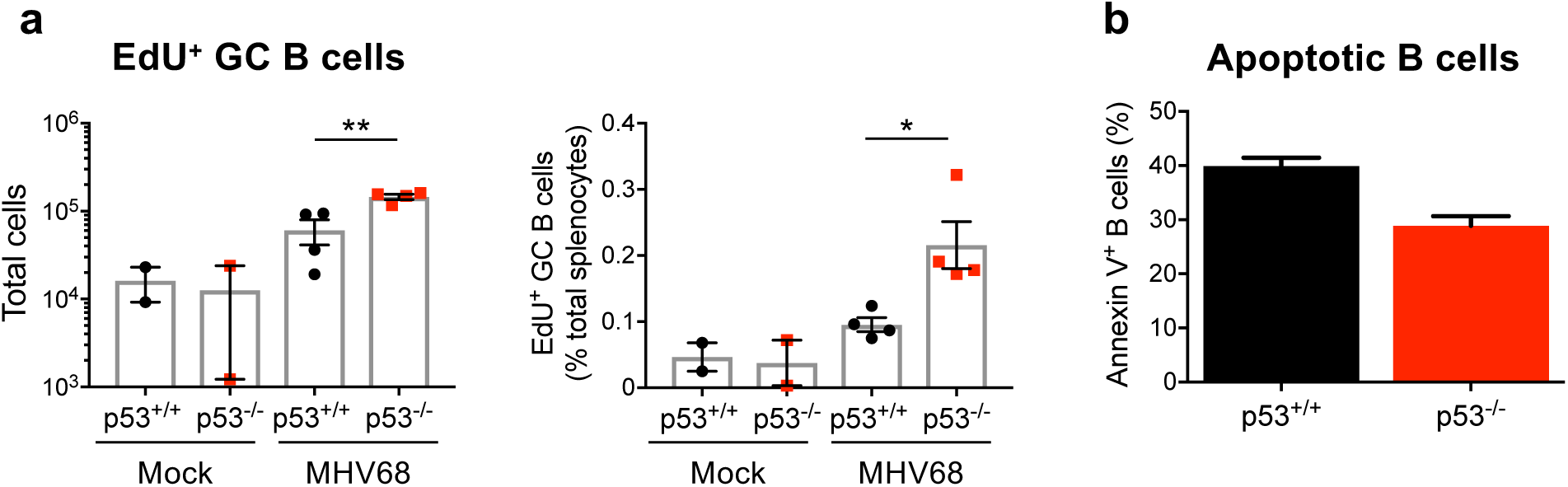
p53 controls GC B cell proliferation during MHV68 latency establishment. **a,** Quantification of GC B cells undergoing DNA replication on day 16 after mock infection or infection of p53^-/-^ and p53^+/+^ mice with MHV68. EdU was injected intraperitoneally 2 hours prior to sacrifice to label newly replicated DNA. Click chemistry was performed and EdU+ GC B cells (gated as CD19^+^/B220^+^/GL7^+^/CD38^lo^) were detected by flow cytometry. Total (left panel) and percent EdU+ GC B cells per spleen (right panel) are shown. **b,** Flow cytometric quantification of B cells (gated as CD19^+^) undergoing apoptosis on day 16 after infection of p53^-/-^ and p53^+/+^ mice with MHV68. Apoptotic cells were identified as Annexin V^+^ after *ex vivo* staining. Gating strategies and additional controls are shown in **Supplemental** Figure 7. Data represent means +/- SEM. Mann-Whitney unpaired *t* test, * p<0.05, ** p<0.005.

### Long-term MHV68 latency is not enhanced in p53-deficient mice

After the initial virus-driven B cell activation and expansion associated with MHV68 infection, both total cell numbers and the frequency of cells harboring viral genomes contract to levels that are relatively stable during long-term infection^61,62,83^. In testing the impact of p53 on long-term MHV68 latency, we found that viral genome-positive cells in spleens of p53^-/-^ mice were ca. 5-fold lower than WT mice on day 42 post-infection, a phenotype that was maintained up to 100 days post-infection (**Fig. S5**). The total number of cells, including total B cells and GC and plasmablast subsets, were equivalent in spleens of both knockout and WT animals by day 42 after infection (**Fig. S5**). These data suggest that p53 is mainly activated to restrict virus-driven B cell expansion during latency establishment and does not play a critical role in controlling infection as latency is maintained over time. Moreover, it is notable that despite restricting initial latent colonization of the host, the modest reduction in latent viral burdens observed at later time-points suggests that p53 induction during latency establishment has a pro-viral role in long-term latent infection by MHV68.

### The M2 latency protein promotes p53 activation in primary B cells

Since p53 was stabilized specifically in MHV68 infected cells where it appeared to restrict GC B cell expansion during early latent infection, we hypothesized that viral proteins involved in latency establishment are responsible for activating p53 in B cells. To test this hypothesis, we performed a screen in which primary B cells were transduced with retroviruses that encode the major MHV68 latency genes, *M2*, *M11*/v-BCL2, *ORF72*/v-Cyclin, and *ORF73*/mLANA^15,16,22,84,85^ and evaluated p53 induction by flow cytometry. Frameshift-stop mutant versions of each gene were used as negative controls. In this screen, only the M2-encoding retrovirus led to p53 stabilization (**Fig. 5a**). As with MHV68 infected splenocytes, only transduced cells exhibited increased p53 detection (**Fig. S8**), which further supports the notion that p53 induction is an intrinsic cellular response to viral gene expression.

**Figure 5:**
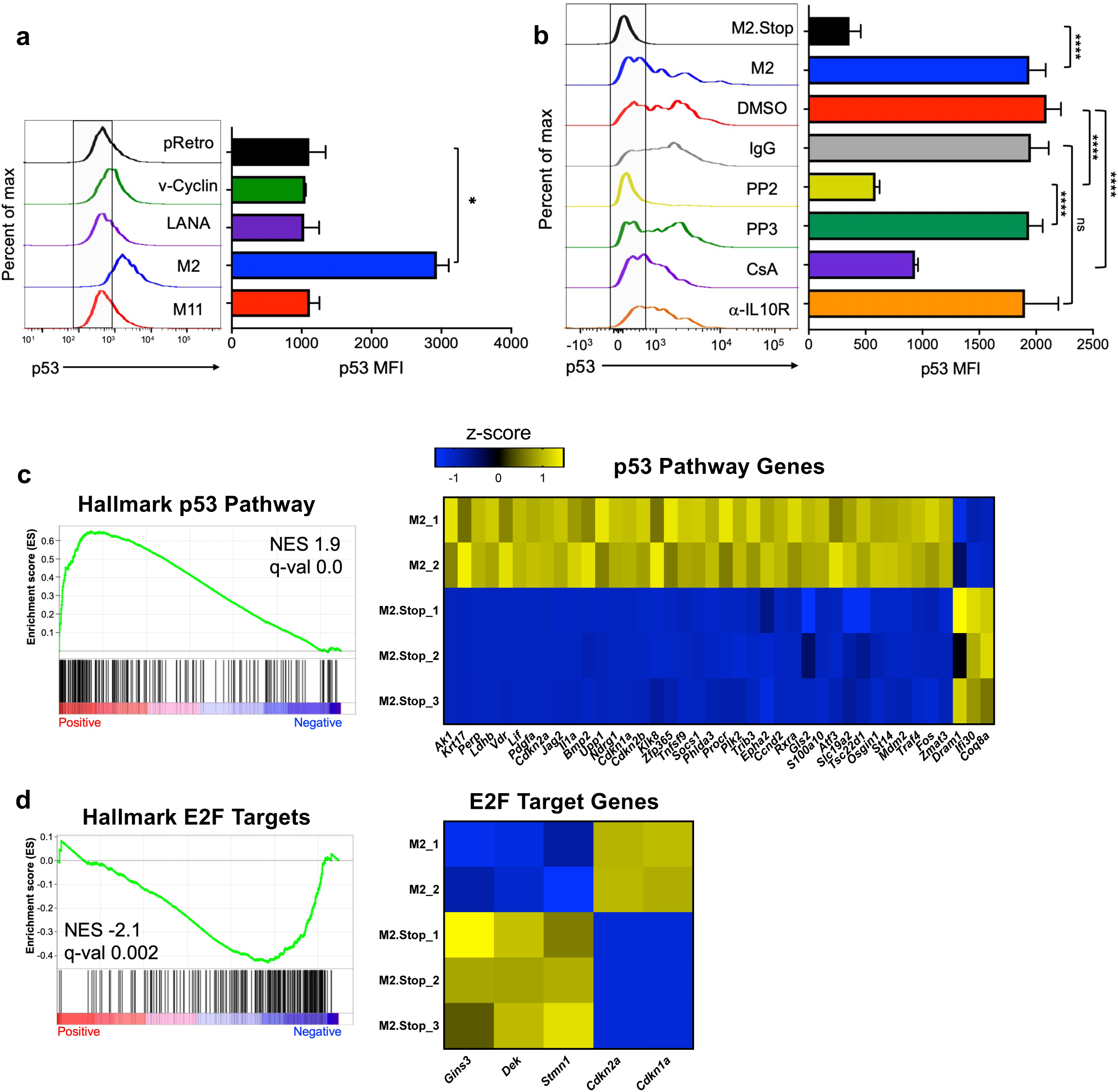
MHV68 latency protein M2 causes p53 induction via Src-family kinase signaling. **a,b,** Flow cytometry analysis of p53 induction in primary murine B cells after transduction with retroviruses that encode MHV68 latency proteins. Representative histograms and mean fluorescence intensities of p53 staining in transduced cells are shown. Data in **a** show p53 staining 4 days after transduction of cells with retroviruses that encode v-Cyclin, LANA, M2, M11, or a frame-shift/translation-stop mutant M2 (M2.Stop) as a negative control. Data in **b** show p53 staining 2 days after transduction with M2.Stop or M2-encoding retroviruses in the presence of DMSO (1:1000), mouse IgG (20 μg/mL), PP2 (10 mM), PP3 (10 mM), CsA (500 ng/mL), or rat anti-IL10R (20 μg/mL). Inhibitors were added 24 hours post-transduction. Gating strategies and additional controls are shown in **Supplemental** Figure 8. Data represent mean +/- SEM. Two-tailed Student’s *t* test, ns - not significant, * p<0.05, ** p < 0.01, and *** p < 0.001. **c-e,** RNA-seq was performed 4 days after transduction of primary B cells with M2 or M2.Stop retroviruses to evaluate changes in transcription. Data in **c** show gene set enrichment analysis (GSEA) and hallmark gene z-score heatmaps comparing M2 to M2.Stop for the hallmark p53 gene set. Data in **d** show the hallmark E2F gene set. The reference list was derived from Hallmark gene sets and compared with a pre-ranked list (by fold) of global average gene expression. Statistical scores are inset into the top right of analysis images. NES, normalized enrichment score, q-val FDR-adjusted *p* value. Heatmaps show average expression data for M2 or M2.Stop expressing primary B cells. The genes presented were derived from GSEA and DEseq2 analysis of all genes with significant expression changes where the expression level increased from M2 to M2.Stop at least 2-fold and decreased from M2 to M2.Stop at least 1.5-fold.

The M2 latency protein of MHV68 functions similarly to EBV latent membrane proteins to promote B cell proliferation and differentiation^57,86^. M2 acts as a scaffold protein that activates Src-family kinases^28,87^, triggering a signaling cascade that results in NFATc1 and IRF4-dependent production of the cytokine IL-10^26^. M2 function is specifically required in GC B cells to facilitate MHV68 latency establishment^30,88^, and its expression in naive B cells results in a GC-like activation profile and BCR class switching that is dependent on IL-10^86^. Given that oncogenes are well known to activate cellular tumor suppressor responses^89^, we reasoned that the M2-driven signal transduction pathway would be recognized as a potentially oncogenic stimulus by the cell, resulting in p53 activity. To test this hypothesis, we transduced primary B cells with control and M2-encoding retroviruses in the presence or absence of specific pharmacologic inhibitors or an IL-10 inhibitory antibody. Remarkably, Src inhibitor PP2 completely blocked M2-mediated p53 induction, while its inactive control analog PP3 had no effect (**Fig. 5b**). Cyclosporin A (CsA), an inhibitor of the calcineurin/NFAT pathway, partially prevented p53 induction, however M2-mediated p53 induction still occurred in the presence of an antibody that blocks the IL-10 receptor. Together and in the context of previously published work^26^, these data strongly suggest that p53 activation is primarily mediated by M2’s activation of a Src-family kinase signal transduction cascade. This is also consistent with Src kinases acting upstream of NFAT and IL-10 upon expression of M2 in B cells.

Detecting p53 induction after expression of M2 in B cells seemed to contradict prior work suggesting that M2 promotes cellular proliferation and survival^26,86,90^. We therefore performed RNA-seq experiments as an unbiased means to evaluate transcription pathways regulated by M2 expression in B cells. In comparisons of control to M2-transduced cells, gene set enrichment analysis (GSEA) revealed a significant positive enrichment of p53 hallmark transcripts upon M2 expression (**Figs. 5c** **and S11**). Consistent with p53 activity restricting cellular proliferation, E2F-related transcripts were downregulated in M2-transduced B cells (**Figs. 5d** **and S11**). Based on these findings, we reasoned that M2 would promote increased proliferation of p53^-/-^ B cells relative to WT cells. Indeed, EdU incorporation assays showed a large increase in the percentage of proliferating p53^-/-^ cells compared to WT cells upon M2 expression (**Fig. 6a**). As in WT cells, EdU incorporation was potently reduced by Src kinase inhibitor PP2 in p53-deficient cells, which further highlights the importance of Src family kinases in M2-mediated B cell activation (**Fig. 6b**). Importantly, RNA-seq analyses revealed that p53 hallmark genes, which include numerous anti-proliferative transcripts, was negatively enriched in p53^-/-^ B cells after M2 transduction (**Fig. 6c** **and S11**), while E2F target genes were induced in the absence of p53. Together, these data demonstrate that M2 drives Src family kinase-related cellular proliferation that is restricted by p53 activation in primary B cells.

**Figure 6:**
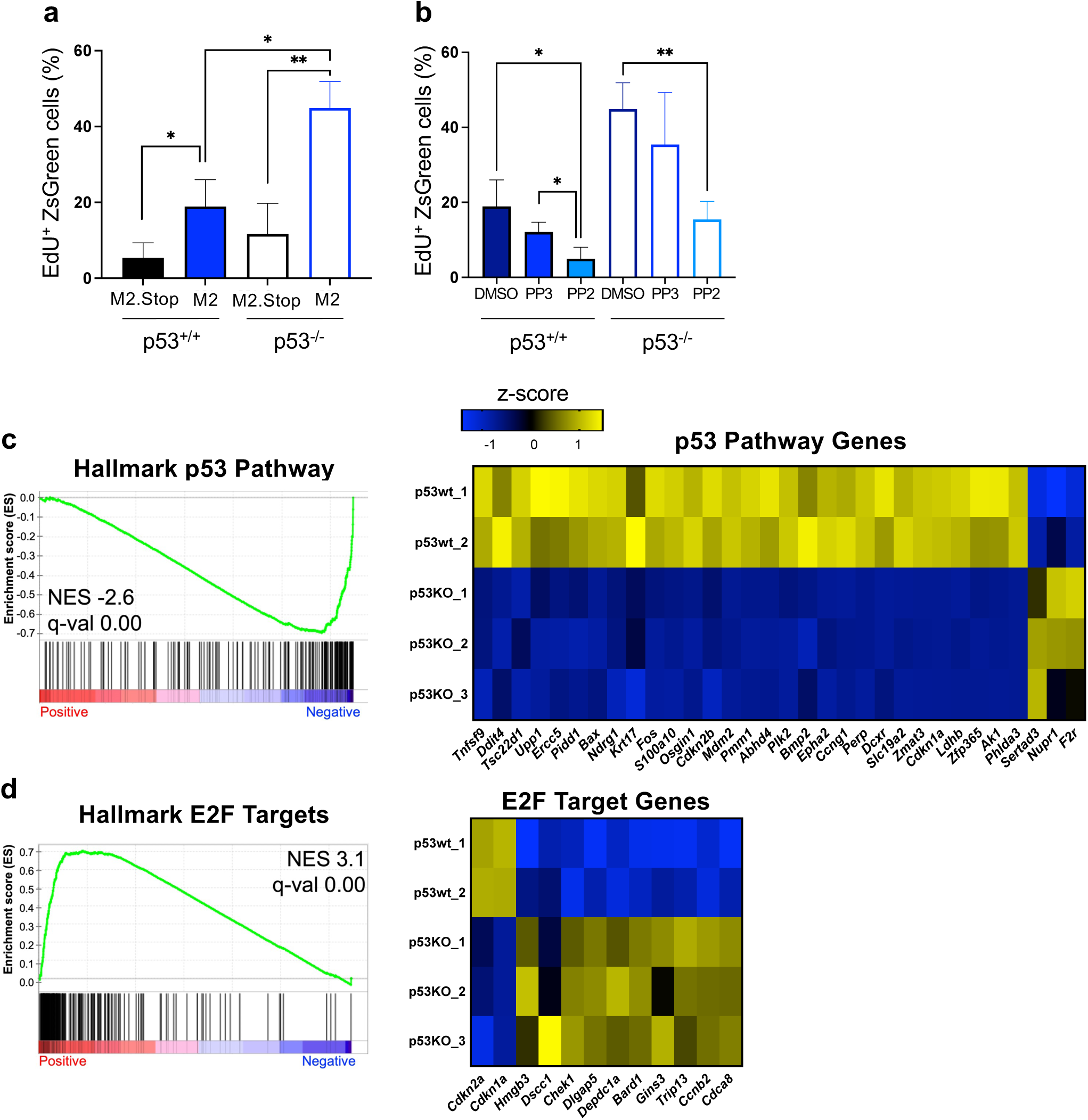
p53 suppresses M2-driven B cell proliferation. **a,b,** Quantification of p53^-/-^ or p53^+/+^ B cells undergoing DNA replication after transduction with M2 or M2.Stop retroviruses. Cells were pulsed with 10 μM EdU for 4 hours prior to harvest at 50 hours post-transduction. Cells in **b** were treated with Src-family inhibitor PP2 (10 mM) or controls (DMSO at 1:1000, PP3 at 10 mM) 24 hours post-transduction. EdU incorporation in newly synthesized DNA was labeled by Click chemistry. EdU+ transduced B cells (gated on ZsGreen+ cells) were detected by flow cytometry. Data represent means +/- SEM. Two-tailed Student’s *t* test, ns - not significant, * p<0.05, ** p < 0.01. Gating strategies and additional controls are shown in **Supplemental** Figure 9. **c-e,** RNA-seq was performed 4 days after transduction of p53^-/-^ or p53^+/+^ B cells with M2 or M2.Stop retroviruses to evaluate changes in transcription. Data in **c** show gene set enrichment analysis (GSEA) and hallmark gene z-score heatmaps comparing M2 to M2.Stop for the hallmark p53 gene set. Data in **d** show the hallmark E2F gene set. Statistical scores are inset into the top right of analysis images. NES, normalized enrichment score. q-val, FDR-adjusted *p* value. Heatmaps show average expression data for M2 or M2.Stop expressing primary B cells. The genes presented were derived from GSEA and DEseq2 analysis of all genes with significant expression changes where the expression level increased from M2 to M2.Stop at least 2-fold and decreased from M2 to M2.Stop at least 1.5-fold.

### EBV LMP1 causes p53 activation in primary mouse B cells

Analogous to MHV68, EBV induces GC-like responses in infected B cells, which the virus usurps to facilitate latency establishment^6,8,91^. In particular, the EBV “growth program” (also known as Latency III) involves the coordinated expression of several latency factors that lead to B cell activation, proliferation, and eventual immortalization into lymphoblastoid cell lines (LCLs)^92,93^. Prior studies demonstrated that EBV infection of primary human B cells elicited p53 stabilization^49^, and p53 agonists limit LCL formation upon EBV infection^51^. We therefore hypothesized that distinct latency factors involved in the EBV growth program may trigger p53 induction, and we focused our analysis on LMP1 due to its functional similarities to M2^57^. Consistent with our hypothesis, transduction of primary mouse B cells with retroviruses encoding LMP1 caused p53 stabilization (**Fig. 7a**). GSEA analyses of transcription in LMP1-transduced cells confirmed that p53-related transcripts were significantly enriched (**Fig. 7c**), as were c-Myc signature transcripts. Comparisons of RNA-seq data from p53 competent and knockout cells expressing LMP1 indicated that enrichment of both c-Myc and p53-related transcription in these cells is linked to p53 expression (**Fig. S11**). Interestingly, both LMP1- and M2-transduced p53^-/-^ B cells formed large clusters of cells (**Fig. 7b**) that proliferated and survived in culture for 2-3 weeks (**Fig. S10**), whereas transduced WT cells could not be maintained. These data demonstrate that EBV-encoded LMP1 elicits p53 activation in mouse B cells in a manner that is consistent with p53 restricting cell survival and proliferation. This parallel observation for an EBV latency protein suggests that antagonism of the virus-driven GC response by p53 is a common feature of GHV latency establishment.

**Figure 7:**
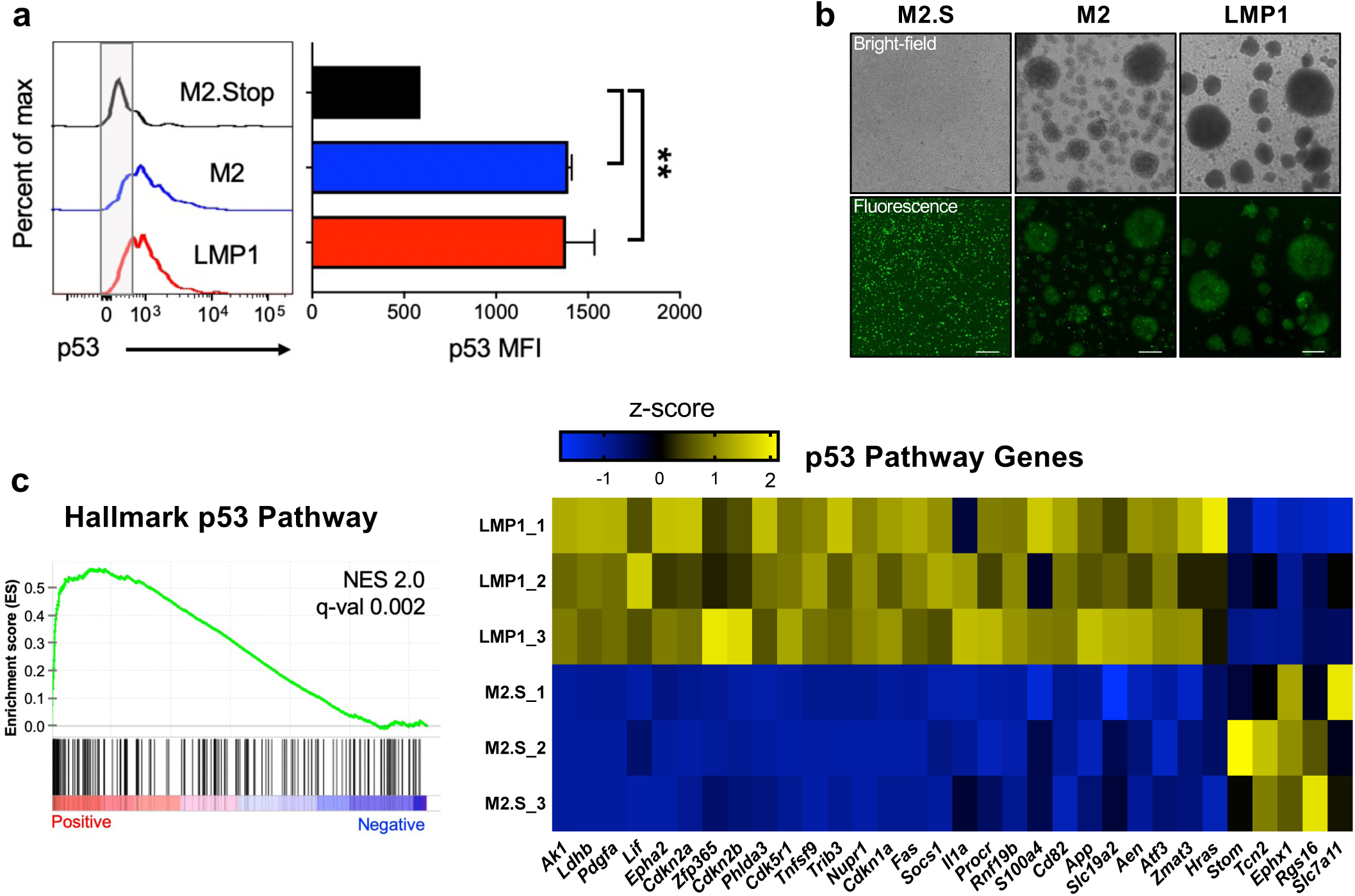
EBV LMP1 activates p53 in mouse B cells. **a,** Flow cytometry analysis of p53 induction in primary murine B cells 4 days after transduction with LMP1, M2 or M2.Stop retroviruses. Representative histograms and mean fluorescence intensities of p53 staining in transduced cells are shown. Data represent means +/- SEM. Two-tailed Student’s *t* test, ** p < 0.01. Gating strategies are shown in **Supplemental** Figure 10. **b,** Transduced B cells were imaged 48 hours post-transduction. Scale bar indicates 250 μm. **c,** RNA-seq was performed 4 days after transduction of primary B cells with LMP1 or M2.Stop (negative control) retroviruses to evaluate changes in transcription. Data show gene set enrichment analysis (GSEA) and hallmark gene z-score heatmaps comparing LMP1 to M2.Stop for the hallmark p53 gene set. The reference list was derived from Hallmark gene sets and compared with a pre-ranked list (by fold) of global average gene expression. Statistical scores are inset into the top right of analysis images. NES, normalized enrichment score. q-val, FDR-adjusted *p* value. Heatmaps show average expression data for LMP1 or M2.Stop expressing primary B cells. The genes presented were derived from GSEA and DEseq2 analysis of all genes with significant expression changes where the expression level increased from LMP1 to M2.Stop at least 2-fold and decreased from LMP1 to M2.Stop at least 1.5-fold. A table of hallmark gene sets with significant changes caused by LMP1 and RNA-seq analysis of LMP1 transduced p53^-/-^ primary murine B cells are shown in **Supplemental** Figure 11.

## DISCUSSION

GHVs take advantage of B cell proliferation and differentiation to facilitate the establishment of life-long latent infection of their hosts. This is accomplished through a coordinated latency gene-expression program that augments and mimics natural developmental processes that guide B cell activation and differentiation as part of the adaptive immune response to infection. We demonstrate here using an *in vivo* model of GHV infection that p53 is induced specifically in infected cells during the period in which a murine GHV, MHV68, manipulates the B cell compartment to colonize its host. Although p53 is thought to be repressed by Bcl-6 in GC B cells to facilitate rapid cellular proliferation and mutagenesis of the B cell receptor^94^, the absence of p53 correlated with large increases in the number of GC B cells, their proliferation, and infection of these cells by MHV68. Further supported by detection of p53-responsive transcripts within latently infected cells, these data strongly suggest that p53 remains responsive in the GC B cell compartment and is capable of restricting GHV-driven B cell expansion as the virus establishes latency. Since p53 stabilization occurs during EBV infection of primary human B cells^49^, p53 agonists restrict their immortalization^51^, and LMP1 induces p53, we propose that p53 is a common intrinsic inhibitor of GHV latency establishment in B cells.

As a master regulator of cell-stress responses, p53 activation during viral infection has been evaluated for numerous viruses, such as influenza virus, herpes simplex virus (HSV), human cytomegalovirus (HCMV), HIV, SV40, adenovirus, and others, including MHV68 and EBV^49,81,95–97^. In fact, p53 was initially identified through its interaction with the SV40 oncoprotein, Large T antigen (T_ag_)^98^. It is important to emphasize that these studies primarily focused on p53 function and inhibition during the productive replication phase of the viral infection cycle. For instance, p53 is potently inhibited by Large T_ag_, adenovirus E1B-55K/E4orf6, and MHV68 mLANA and muSOX during lytic replication^96,99^. Identical viral replication in both WT and p53^-/-^ mice highlights the potent inhibition of p53 during acute MHV68 replication. In contrast, p53 is an interferon stimulated gene that participates in the antiviral response and host survival during HSV-1 infection^100^, whereas HCMV integrates p53 activity to promote lytic gene expression and viral replication^97^. While well-studied in productive replication of numerous and diverse virus infections, surprisingly little is known regarding p53 function in the latent phase of herpesvirus infection. Certainly, prior studies suggest that some GHV latency proteins inhibit p53 (summarized below), but whether interactions between a GHV and p53 impact latency in a living host was previously unexplored. To the best of our knowledge, the data presented here represent the first *in vivo* evidence of an antagonistic relationship between host p53 and the GHV latency-associated transcription program that influences viral latency outcomes.

Since the absence of p53 correlated with increased GC proliferation and early viral latency establishment, our data strongly suggest that p53 is not simply inhibited by viral latency factors, an idea that has long been proposed in the GHV field, but that has become somewhat controversial in recent years. For instance, EBV nuclear antigen-1 (EBNA-1), a DNA-binding protein that facilitates maintenance of the latent viral episome, is expressed broadly in EBV cancers and hypothesized to facilitate cellular immortalization by promoting p53 proteolysis via an interaction with ubiquitin-specific protease 7 (USP7)^54,101^. However, despite expression of EBNA-1 and in agreement with our findings, p53 is activated early during EBV infection of primary B cells, and LCLs remain responsive to p53 agonists^49,51^. Although the KSHV episome maintenance protein LANA inhibits p53 in overexpression systems^52,102,103^, p53 remains functional in PEL cells that contain high levels of LANA protein^55,104^. While p53 can be inhibited by the MHV68 LANA homolog, the inhibitory function appears to be restricted to the lytic replication cycle, where it requires an undefined activating event and is assisted by additional lytic-phase viral proteins^59,96^.

Given that inactivation of p53 by mutation or suppression is predisposing to cellular transformation, it is perhaps not surprising that viruses that establish lifelong chronic infections as part of their maintenance strategy would evolve to *not* block p53 function within latently infected cells. Presumably, potent inhibition of p53 would reduce host survival and limit viral transmission. However, it is important to also emphasize that this idea does not rule out the possibility that p53 responses are toned down by viral latency factors either transiently or subtly to enable GHVs to overcome p53-dependent barriers to latency establishment during initial colonization of host lymphoid tissues. It is additionally notable that p53 is frequently mutated in eBL and PEL^2^, suggesting that p53 function is a critical determinant of pathogenesis for these diseases. p53 knockout mice are primarily predisposed to thymic lymphoma and sarcomas^105^, whereas people with p53 mutations exhibit a broad spectrum of cancers^106^. Whether GHV infection influences the tumor phenotype of hosts when p53 is non-functional is unknown but will be of interest to determine using the mouse model moving forward, as we suspect that enhanced B cell proliferation during MHV68 infection of p53-deficient animals is a hallmark of incipient oncogenesis.

With regard to a specific mechanism for p53 induction during GHV latency establishment, increased GC expansion during infection of p53 knockout mice supported a model in which viral latency proteins that drive B cell proliferation and differentiation lead to the activation of p53. In agreement with known responses to viral oncogene expression^56,80,81^, we suspected that p53 would then serve to counteract the proliferative expansion of latently infected cells promoted by the viral latency program. Indeed, identification of the latency-specific gene product M2, which in-and-of-itself promotes GC B cell expansion^30,86^, as a potent p53 activator in primary B cells directly supports this hypothesis. M2 serves as a molecular scaffolding protein that binds and activates Src-family kinases^27,28,87^. This leads to induction of NFAT/calcineurin signaling pathways that promote IL-10 production and B cell differentiation^26^. We demonstrated that signal transduction via Src-family kinases is the key event responsible for triggering p53 in response to M2 expression, and not IL-10-related effects on the cell. It is well established that aberrant cellular activation and cell-cycle progression via oncogenes such as H-Ras and c-Myc triggers tumor suppressor responses, especially activation of p53-dependent inhibition of cell-cycle progression^107,108^. Our data therefore support the conclusion that M2/Src activity is recognized by an infected B cell as a potential oncogenic threat, leading to p53 activity that restricts infected cell proliferation.

It will be of interest to determine if MHV68, like EBV^37,109,110^, elicits ATM and/or ATM and Rad3-related (ATR) DNA damage responses as a mechanism of p53 phosphorylation and stabilization or if other kinases fulfill this role. For EBV, ATM and ATR are activated in response to a period of hyperproliferation and replication stress and limit the immortalization of infected B cells in culture^37,111,112^. The functions of ATM in MHV68 infection of mice have been evaluated, and MHV68’s relationship with ATM is complex in that ATM expression in B cells does not restrict, but rather facilitates viral latency establishment^43,45^. One wonders if the reduction in MHV68 latency observed at late timepoints during infection of p53^-/-^ mice is related to the ATM knockout phenotype, perhaps reflecting the accumulation of DNA damage and subsequent death of the infected cells. Of note, we do find evidence of increased genomic instability in p53^-/-^ mice after MHV68 infection (S.M. Owens and J.C. Forrest, manuscript in preparation). A specific role for ATR, which is primarily activated by single-strand DNA breaks that occur as a result of DNA replication stress^113^, would support the notion that aberrant cell-cycle progression and DNA replication caused by M2/Src is a trigger for p53 activation. In further support of this notion, a previous RNA-seq analysis of MHV68-infected splenocytes revealed enhanced transcription of genes involved in the ATR signaling pathway, in addition to p53-related transcription^114^. While the ATR hypothesis is compelling, dissecting this pathway *in vivo* would be challenging, as ATR-deficient mice exhibit a progeria syndrome and immune dysfunction^115^.

Although M2 is unique to MHV68, it exhibits functional similarities to latency proteins encoded by human GHVs, such as EBV LMP1/LMP2A and KSHV K1/K15, especially integration of signaling events that occur upon B cell receptor and co-stimulatory molecule crosslinking^57,91,116–119^. Combined expression of EBV LMP1 and LMP2A in a transgenic mouse model caused GC B cell expansion and B-cell differentiation into rapidly growing plasmablasts that have enriched expression of E2F targets and G2M checkpoint genes^120^, which is analogous to M2 effects on B cells^86^. Both LMP1 and LMP2A mimic aspects of B cell signaling, but only LMP1 is a *bona fide* oncogene^36,40,121,122^, which motivated our testing of LMP1-mediated p53 induction. Depending on the system being tested, LMP1 is reported to both induce p53-stabilization and cell-cycle inhibition or conversely to prevent p53 function^123,124^. Our data indicate that a p53 anti-proliferative response is activated in response to LMP1 where it prevents the outgrowth of transduced B cells. Interestingly, p53 stabilization was enhanced in lymphomas that develop in a transgenic mouse model in which LMP1 is constitutively expressed in B cells^122^. In light of our data, we suspect that LMP1-driven B cell activation triggers p53 in an attempt to control tumor cell growth. While the percentage of cells with stabilized p53 was not evaluated in the transgenic mouse model, it is interesting to note that tumors were also characterized by NF-kB activation, which is necessary to overcome p53-mediated growth restriction during LCL formation^51^. Moving forward, it will be of interest to determine if LMP2A, K1 and K15 (and other human GHV latency proteins) also activate p53. Since LMP2A reduces LMP1 hyperactivation of B cells^125^ and combined LMP1/LMP2A-deficient EBV exhibits diminished transforming capacity in a cord blood model of EBV lymphomagenesis^126^, it will also be important to investigate whether LMP2A suppresses p53 induction by LMP1.

In total, our data suggest a dynamic between p53 and viral latency genes that is analogous to an automobile’s gas pedal and brake. The cellular activating functions of the viral latency genes are necessary for the latently infected cell to progress from point A (initial infection of a naïve B cell) to point B (progression through the GC reaction to enable long-term latency in memory B cells or reactivation from plasma cells). Without the braking mechanism provided by p53, the process would careen out of control, leading to either a reduction in overall latency (**Fig. S5**) or lymphoma. While roles for p53 in preventing aberrant cell-cycle progression, maintaining genomic integrity, and limiting cancer development are well defined, the impact of p53 on pathogenic outcomes in GHV-infected B cells remains to be determined. Since p53^-/-^ B cells transduced with either M2 or LMP1 proliferated and could be maintained in culture, whereas transduced WT cells could not, we hypothesize that the absence of p53 will hasten B cell transformation when these latency factors are expressed. In addition to evaluating M2 and LMP1-mediated cellular transformation, experiments are underway to determine whether infection with MHV68 alters the tumor spectrum of p53 knockout mice. Given the increase in GC B cell proliferation, we are particularly excited to define the impact of MHV68 infection on B cell lymphomagenesis. There is an ongoing need for *in vivo* models of GHV infection-associated lymphoma development, and these experiments could provide new models for defining GHV drivers of oncogenesis and pathways for targeting therapeutically.

## MATERIALS AND METHODS

### Cell culture and viruses

Swiss albino 3T3 fibroblasts were purchased from ATCC. Murine embryonic fibroblasts (MEFs) were harvested from C57BL/6 mice embryos and immortalized as previously described^73^. All cells were cultured in DMEM supplemented with 10% fetal bovine serum, 2 mM L-glutamine, and 100 U/ml penicillin/streptomycin. Cells were maintained at 37°C in 5% CO_2_. Viruses used in this study were previously described and include WT MHV68^127^, H2B-yellow fluorescent protein (YFP)-expressing MHV68^60^, Cre-recombinase-expressing MHV68^65^, and mLANA-null MHV68 (73.STOP)^16^. Viral stocks were generated as previously described^59^. Viral titers were determined by MHV68 plaque assay^128^.

### Mice and infections

All mice were housed and cared for according to the guidelines of UAMS Department of Laboratory Animal Medicine and all state and federal requirements. p53-null mice on a C57BL/6 background (B6.129S2-*Trp53^tm1Tyj^/*J) were purchased from Jackson Laboratories and bred p53^+/-^ x p53^+/-^ to get a distribution of p53^+/+^, p53^+/-^, and p53^-/-^ progeny. 7–11-week-old mice were infected with 10^4^ PFU of WT MHV68, H2B-YFP MHV68, MHV68-Cre, or mLANA-null MHV68 by intranasal inoculation or by intraperitoneal injection. Mice were sacrificed according to normal endpoint protocols at the first sign of illness or tumor development. Blood was collected on days 0, 16, and 42 post-infection via the submandibular vein.

### Splenocyte isolation and limiting-dilution analyses

Spleens were homogenized in a tenBroek tissue disrupter. Red blood cells were lysed by incubating tissue homogenate in 8.3 g/L ammonium chloride for 10 minutes at room temperature with shaking. Cells were filtered through a 40-micron mesh to reduce clumping. Frequencies of cells harboring MHV68 genomes were determined using a limiting-dilution, nested PCR analysis as previously described^73^. Frequencies of latently-infected cells capable of reactivating were determined using a limiting-dilution analysis for cytopathic effect induced on an indicator MEF monolayer as previously described^73^.

### Antibodies, tetramers, and treatments

CD19-BV650 (6D5), IgM-BV421 (RMM-1), CD38-Pacific Blue (90), and purified B220 (RA3-6B2) were purchased from Biolegend (San Diego, CA). CD3e-PCPCy5.5 (145-2c11), CD8α-PCPCy5.5 (53-6.7), CD4-AF700 (RMA-5), B220-AF700 (RA3-6B2), GL7-eF660 (GL-7), CD38-PE-Cy7 (90), CD138-BV711 (281-2), NK1.1-PCPCy5.5 (PK136), and IgD-PE (11-26c) were purchased from eBioscience. Other antibodies include mouse anti-p53 Alexa Fluor 647-conjugate (Cell Signaling), goat anti-GFP to enhance YFP detection (Rockland), and donkey anti-goat Alexa Fluor 488-conjugate (Invitrogen). MHV68-specific MHC class I tetramers were generated by the NIH Tetramer Core. As a positive control for p53 induction, bulk splenocytes were treated with 10 grays of gamma radiation by exposure to a cesium-137 source. Following exposure, cells were allowed to recover at 37°C in a tissue-culture incubator for 1 hour prior to staining for flow cytometry. Annexin V-PE and PI were added along with antibodies to evaluate cell death.

### Flow cytometry

Cells were washed with FACS buffer (0.2% BSA, 1 mM in PBS) before blocking with Fc block (Invitrogen) and incubation with eF780 live/dead viability stain (eBioscience) for 10 minutes at 4°C. Surface staining was then performed with antibodies diluted at 1:300 for 30 minutes incubation time at 4°C. For intracellular stains, cells were fixed and permeabilized using a FoxP3 staining kit (eBioscience) following the manufacturer’s guidelines. The data were collected using an LSRFortessa (Becton Dickinson) and analyzed using FlowJo (10.4.2) software.

### Immunohistology and microscopy

Tissues were fixed in 4% formaldehyde prior to paraffin embedding and cut into 5 µm sections. For H&E, sections were stained with hematoxylin and eosin as previously described^129^. Tissue sections were imaged on an EVOS digital microscope with 10X or 40X objectives (Thermo). For immunohistochemical analyses, the sections were deparaffinized, rehydrated, and incubated in citrate buffer for antigen retrieval before staining with Vectastain Elite ABC kit (Vector) using a chromogenic reporter, DAB (Dako). Sections were counterstained with hematoxylin. Anti-fade mounting medium (Vector) was used prior to examining tissue sections on an Eclipse T5100 microscope (Nikon).

### Real-time PCR and genotyping

Lungs were harvested from mice infected with H2B-YFP MHV68 on day 7 post-infection and homogenized using 0.5 mm silica beads in a Minibeadbeater (BioSpec) as previously described^73^. DNA was extracted from homogenized tissue using a DNeasy kit (Qiagen) according to the manufacturer’s instructions followed by treatment with RNAse I (Qiagen) for 30 minutes at 37°C. Quantitative PCR was performed on 500 ng of DNA with primers forward corresponding to the MHV68 ORF59 genomic locus (forward: 5’-ATG-CAG-ACC-TTC-CAG-CTT-GAC-3’; reverse: 5’-CTC-TTC-CAA-GGG-AGC-TTG-CG-3’). Cycling parameters were 95°C for 30 seconds, 55°C for 30 seconds, and then 72°C for 30 seconds for 40 cycles on an Applied Biosystems StepOnePlus PCR system. Genotyping of mice was performed using a mouse genotyping kit (Kapa Biosystems). Briefly, DNA was extracted from mouse tail snips and amplified as previously described^105^.

### Reverse transcriptase PCR

Splenocytes were isolated from H2B-YFP MHV68-infected or mock-infected mice at 16 days post-infection, and total RNA was extracted from the cells using Qiagen RNeasy Mini Kit (#74104). 1 ug RNA was used for cDNA synthesis (Invitrogen) prior to PCR analysis following manufacturer’s guidelines (Agilent). Gene-specific TaqMan probes (Applied Biosystems) for Mdm2 (Mm01233136_m1), Cdkn1a (Mm00432448_m1), and β-actin (Mm00607939_s1) were utilized for quantification. Reactions were performed in an Applied Biosystems StepOnePlus PCR system with cycling conditions of 10 minutes at 95°C followed by 40 cycles of 15 seconds at 95°C and 1 minute at 60°C. Biological triplicate samples were analyzed in technical triplicate using the ΔΔCT method with β-actin as the cellular housekeeping transcript control.

### P53 pathway transcriptional array

Ai6 mice were infected with 10^4^ PFU of MHV68-Cre intranasal inoculation and splenocytes were isolated at 16 days post-infection. One thousand ZsGreen positive and negative splenocytes were sorted in biological triplicate to a 96 well plate using a FACSAria II (BD). Transcriptome amplification was performed using QIASeq Stranded RNA library kit (Qiagen). RT^2^ Profiler PCR Array Mouse p53 Signaling Pathway (PAMM-027Z – Qiagen) was used for quantification. Reactions were performed in an Applied Biosystems StepOnePlus PCR system with cycling conditions of 10 minutes at 95°C followed by 40 cycles of 15 seconds at 95°C and 1 minute at 60°C. Biological triplicate samples were analyzed using the ΔΔCT method with β-actin as the cellular housekeeping transcript control.

### Plasmids

Retrovirus plasmids were generated by restriction enzyme cloning of latency genes into pRetroX-IRES-ZsGreen (Takara). Latency locus genes were amplified from WT MHV68-BAC DNA^127^ and the frame-shift stop mutant, M2.Stop, was amplified from M2.Stop MHV68-BAC DNA^130^ using primers in Table 2. LANA was cloned into the pRetroX vector via SacII and BamHI. vCyclin, M2, M11, and M2.Stop were amplified and cloned via NotI and BamHI. The LMP1 coding region was amplified from MSCV-N LMP1^131^ (Addgene plasmid #37962) and cloned into pRetroX-IRES-ZsGreen via Gibson Assembly. All plasmids were confirmed by sequencing (Plasmidsaurus).

**Table 1.**
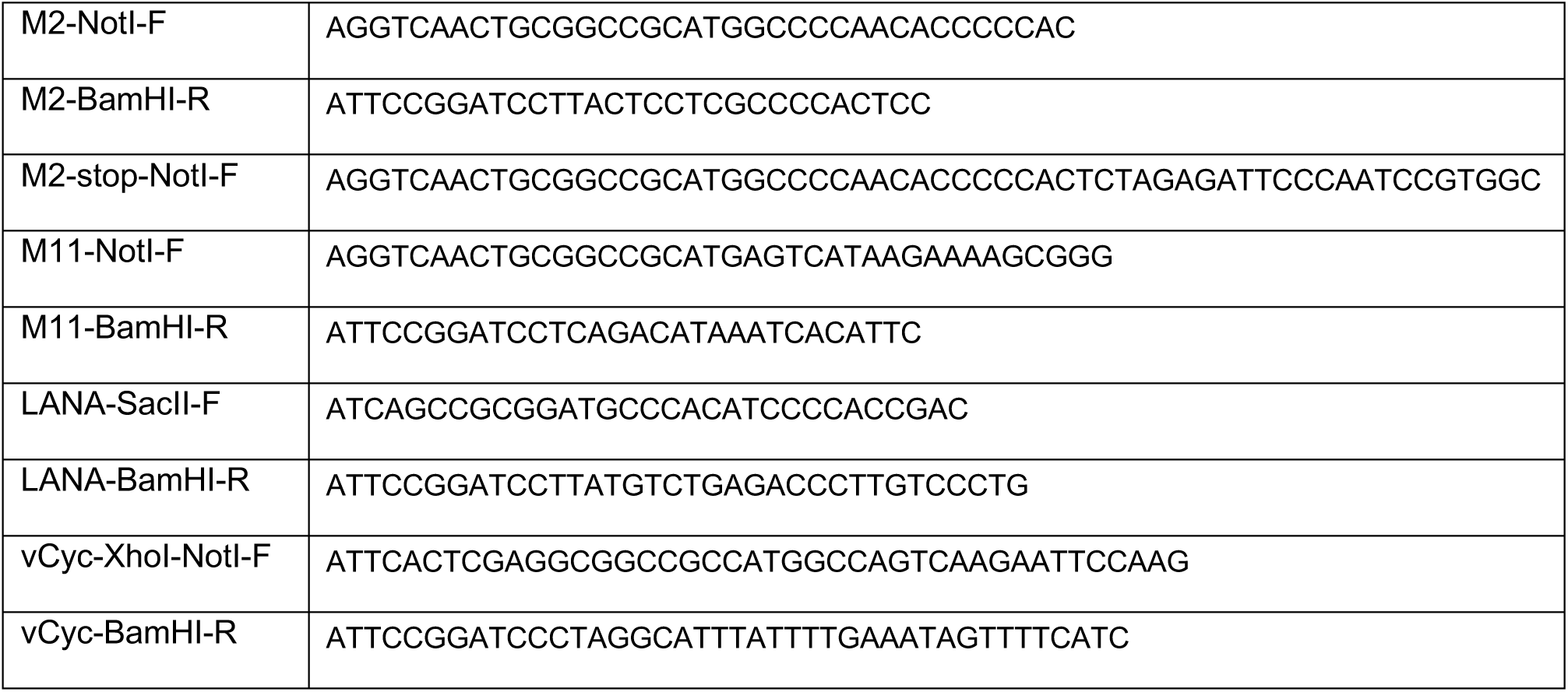
pRetro_IRES_ZsGreen cloning primers.

### B cell cultures and retroviral transduction

Naïve B cells were isolated from spleens by immunomagnetic depletion using MojoSort Pan B cell Isolation Kit (Biolegend) and cultured in RPMI supplemented with L-glutamine, sodium pyruvate, penicillin-streptomycin, 50 μM 2-mercaptoethanol, and 10% fetal bovine serum. B cells were stimulated with 25 μg/ml LPS (Sigma). Retroviral supernatants were obtained from Phoenix cells (ATCC) 72 hours after transient transfection (Lipofectamine LTX, ThermoFisher) with the retroviral plasmids and filtered through a 0.45 mm syringe filter. B cells were transduced 24 hours after LPS stimulation with retroviral supernatants by centrifugation for 30 mins at 1000xg at 37°C in the presence of 5μg/ml polybrene. Cells were washed and supernatants were replaced with RPMI. Triplicate wells per condition were analyzed by flow cytometry at day 4 post-transduction.

### Drug Treatments

Cyclosporin A (500 ng/mL, TCI America), PP2 (10 µM, Selleck Chemical LLC) and PP3 (10 µM, Tocris Bioscience), VE-821 (1 μM, Tocris Bioscience), and KU-55933 (5 μM, Tocris Bioscience) were reconstituted in DMSO. For antibody treatments, mouse IgG Isotype Control (20 μg/mL, ThermoFisher) and anti-mouse IL-10R (20 μg/mL, BioXCell) were diluted in cRPMI. The drugs were used at a final concentration as indicated in parentheses above.

### EdU incorporation assays

Mice were injected with 100 μg of 5-ethynyl-2’deoxyuridine (EdU, Invitrogen) 2 hours prior to splenocyte harvest. Cells were fixed with 4% paraformaldehyde in PBS and Click-iT chemistry (BD Pharmagen) was performed according to the manufacturer’s instructions to fluorescently mark EdU+ cells.

### Enzyme-linked immunosorbent assays

Viral antigen was prepared by infecting 3T12 fibroblasts at an MOI of 0.5 PFU/cell for 96 hours. Cells were washed and fixed in 1% PFA. Viral antigen was used to coat high-binding plates (Denville) overnight at 4°C. After blocking in 5% FBS in PBS, serially diluted serum was incubated on the plate, followed by incubation with HRP-conjugated IgM or IgG Ab (Southern Biotech). SureBlue substrate (KPL) was added to detect the Ag-specific Abs. The wells were read on a FLUOstar Omega plate reader (BMG Labtech).

### RNAseq

Transduced B cells were collected 4 days post-transduction. RNA was harvested using Trizol. The library was constructed by BGI Laboratories using DNBSEQ Eukaryotic transcriptome protocol. Following mRNA isolation, the mRNA was fragmented, and first-strand synthesis performed. Double-stranded cDNA fragments were subjected to end-repair, and a single ’A’ nucleotide was added to the 3’ end of the blunt fragment. Following library QC, single-strand cDNAs were circularized and replicated via rolling cycle amplification. DNA nanoballs were generated and loaded into patterned nanoarrays using high-intensity DNA nanochip technique and sequenced through combinatorial Probe-Anchor Synthesis (cPAS). Sequencing was performed on a DNBSEQ-G400 (BGI Genomics Co). Raw sequence reads were quality-checked using fastQC and trimmed based on quality (Phred >30). After trimming, reads <30 bp in size were discarded. Trimmed sequence reads were mapped to reference genome mm10 using STAR aligner with default parameters^132^. The gene count tables were extracted from alignment results using bedtool2 software^133^. Read counts were normalized using Voom method^134^. Normalized read counts were analyzed using DEseq2^135^. Representative gene set enrichment analysis (GSEA)^136,137^ was performed using the reference list derived from Hallmark gene sets^138^ and compared with a pre-ranked list (by fold) of global average gene expression.

### Data Accessibility

mRNAseq data are deposited under GEO Accession # GSE225579 (Reviewer token: wnwdoamcrbupnat). (https://www.ncbi.nlm.nih.gov/geo/query/acc.cgi?acc=GSE225579).

## Acknowledgments

We thank Andrea Harris in the UAMS Flow Cytometry Core. This work was supported in part by grant R01 CA167065 from the NIH National Cancer Institute and start-up funds from the UAMS College of Medicine and Arkansas Biosciences Institute to J.C.F. The Flow Cytometry Core and the work described here were also supported in part by the Center for Microbial Pathogenesis and Host Inflammatory Responses award P20 GM103625 from the NIH National Institute of General Medical Sciences Centers of Biomedical Research Excellence. S.M.O. was supported by Translational Research Institute (TRI) grant TL1 TR003109 through the NIH National Center for Advancing Translational Sciences. M.M. is supported by K22 CA241355 from the National Cancer Institute with additional support from P30 GM145393 from the National Institute of General Medical Sciences. The funders had no role in study design, data collection, and interpretation, or the decision to submit the work for publication.

## Author’s Contributions

S.M.O., J.M.S., and J.C.F designed the study, analyzed data, and wrote the manuscript. G.L. constructed and tested retroviral vectors. S.J.M. analyzed data and wrote the manuscript. E.S. participated in the study design. M.M. performed RNA-seq analyses and wrote the manuscript. D.G. and J.S. assisted with flow cytometry data analysis. J.C.F. acquired funding. All authors read and approved the final manuscript.

