## Supplemental for "Intrinsic p53 Activation Restricts Gammaherpesvirus-Driven Germinal Center B Cell Expansion during Latency Establishment"

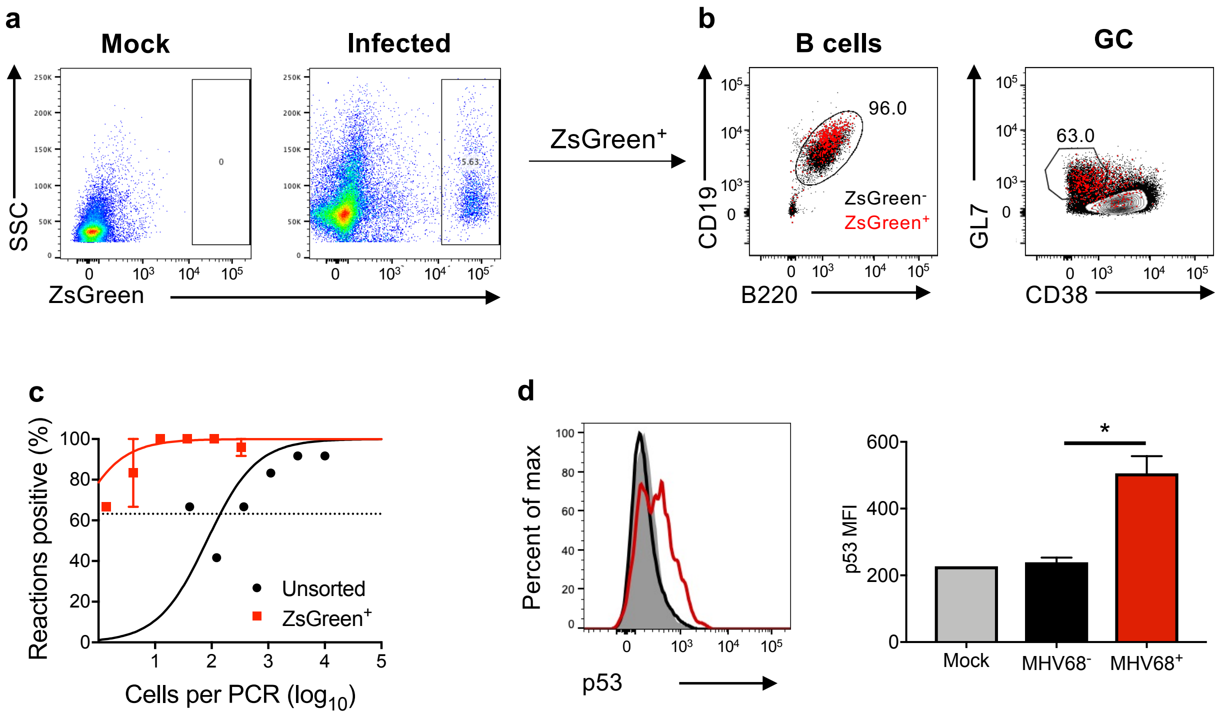


**Supplemental Figure 1: Cre-loxP reporter mouse system identifies MHV68-infected B cells.**
**a-d,** Cre-reporter mice, Ai6 (Zsgreen) or Ai14 (tdTomato), were infected intranasally with MHV68-Cre ($10^4$ PFU). Mediastinal lymph nodes (MLN) and splenocytes were harvested on day 16 post-infection. **a,** Representative flow plots for ZsGreen expression in splenocytes. **b,** Harvested splenocytes were stained with cell-specific markers for the indicated cell types. B cells were gated as $CD19^+/B220^+$, GC B cells as $GL7^+/CD38^{lo}$. $ZsGreen^+$ B cell populations indicated in red, $ZsGreen^-$ B cell populations indicated in black. Population frequencies apply to $ZsGreen^+$ cells. **c,** $ZsGreen^+$ cells were sorted, and limiting dilution PCR was performed to quantify the proportion of splenocytes harboring latent viral genomes. **d,** Cells from the MLN were stained with B cell markers ($B220^+/CD19^+$) and p53. Flow cytometry was performed to evaluate p53 expression in mock-infected (gray), $MHV68^-$ ($tdTomato^-$), and $MHV68^+$ ($tdTomato^+$) B cells. The mean fluorescence intensity of p53 is enumerated in right panel. Two-tailed Student's *t* test, * $p<0.05$.

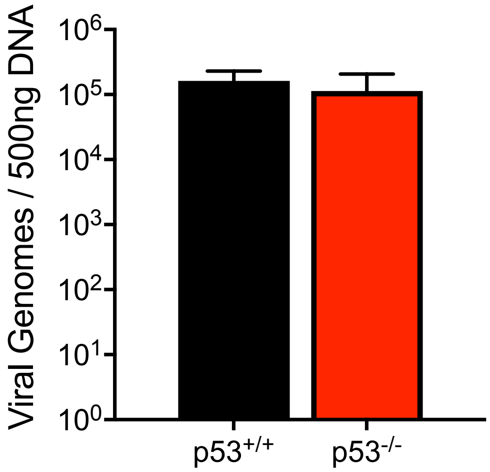


**Supplemental Figure 2: p53 does not affect MHV68 acute replication.** $p53^{+/+}$ or $p53^{-/-}$ mice (n=3) were intranasally inoculated with $10^4$ PFU of H2B-YFP-expressing MHV68. Animals were sacrificed 7 days post-infection and DNA was isolated from lungs for quantitative PCR to detect viral genomes. Results are means of 3 samples +/- SD.

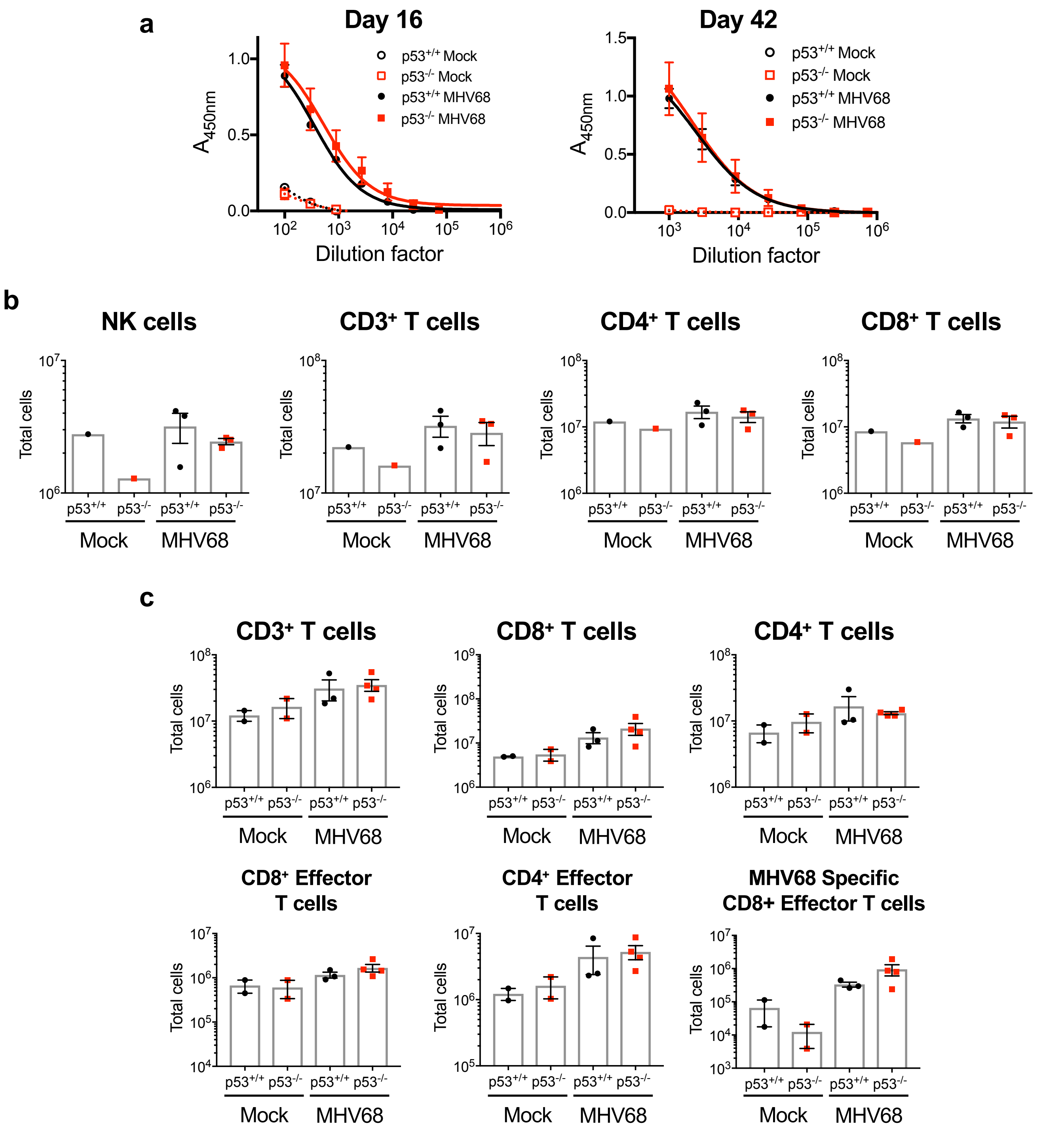


**Supplemental Figure 3: p53 is not required for virus-specific adaptive immunity. a-c,** p53$^{+/+}$ or p53$^{-/-}$ mice were intranasally inoculated with $10^4$ PFU of H2B-YFP-expressing MHV68. **a,** Serum was collected on days 16 and 42 post-infection. Serial dilutions were evaluated by ELISA to quantify MHV68-specific IgG. **b,** Mice were sacrificed on day 16 post-infection and splenocyte populations were analyzed by flow cytometry. Total T cells were gated as CD3$^+$ and specific subsets were identified by CD8 or CD4 expression. Natural killer cells were gated as CD3$^{lo}$/NK1.1$^+$. Live cells were identified with eFluor 780 viability dye. **c,** Mock-infected and MHV68-infected mice were sacrificed on day 42 post-infection. Flow cytometry was performed to quantify total T cells (CD19$^-$/CD3$^+$), CD8$^+$ T cells, CD4$^+$ T cells, effector T cells (CD62L$^-$/CD44$^+$), and MHV68-specific effector CD8 T cells that were detected by staining with an MHV68 ORF6 MHCI tetramer. Data from two independent experiments are shown. Data represent means +/- SEM. No significant differences were present by Mann-Whitney unpaired *t* test.

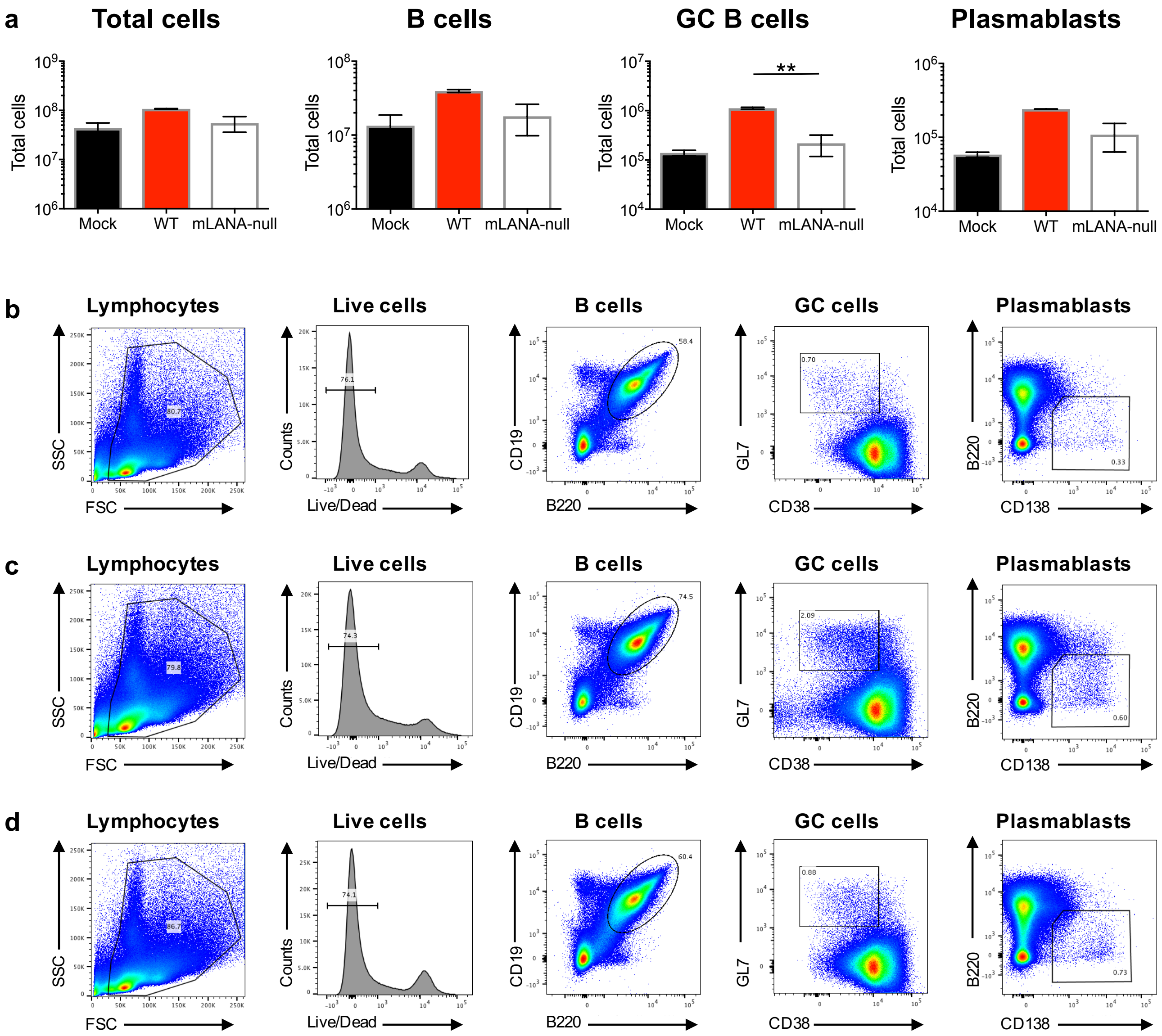


**Supplemental Figure 4: Latent MHV68 infection is required for splenic B cell expansion in p53$^{-/-}$ mice. a-d,** p53$^{-/-}$ mice were mock infected or infected intranasally with $10^4$ PFU of H2B-YFP MHV68 or mLANA-null MHV68 (n=3). Mice were sacrificed on day 16 post-infection and splenocyte populations were defined by flow cytometry. Representative flow cytometry plots are shown for mock infection in **a**, H2B-YFP MHV68 infection in **b**, and mLANA-null MHV68 infection in **c**. B cells were gated as CD19$^+$/B220$^+$, GC B cells as GL7$^+$/CD38$^{lo}$ subset of B cell gate, and plasmablasts as CD138$^+$/B220$^{lo}$. Live cells were identified with eFluor 780 viability dye. Data in **d** represent means +/- SEM. Mann-Whitney unpaired *t* test, ** p<0.01

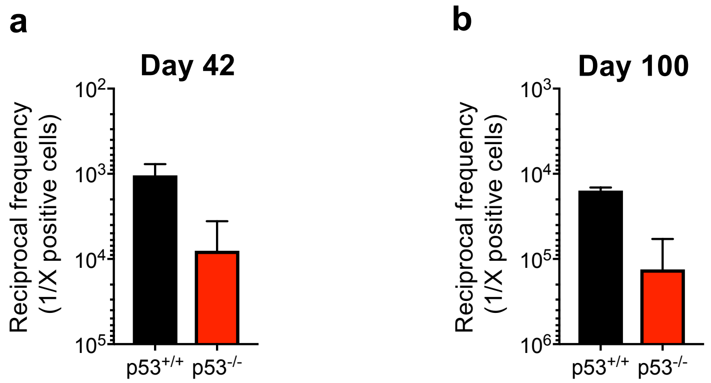


**c**

| | 1/X cells infected | |
|---|---|---|
| **Genotype** | **Day 42** | **Day 100** |
| **$p53^{+/+}$** | 1050 | 6200 |
| **$p53^{-/-}$** | 16000 | 93000 |

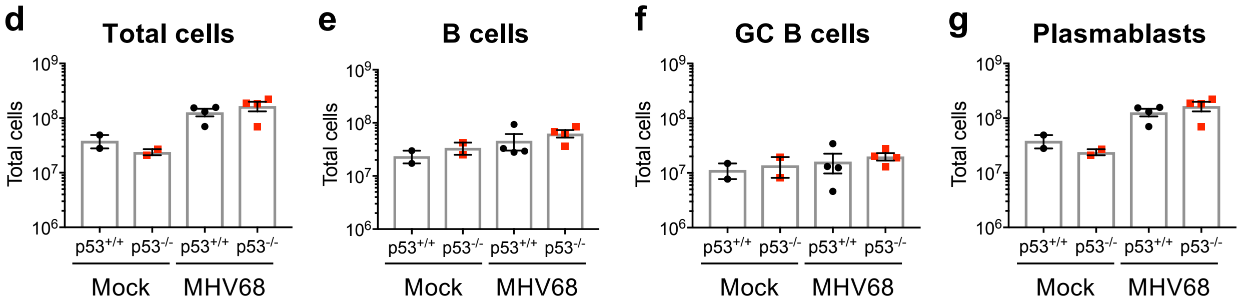


**Supplemental Figure 5: Long-term MHV68 latency is reduced in p53-deficient mice. a,b,** A limiting-dilution PCR analysis was performed to determine the number of splenocytes harboring MHV68 genomes on day 42 (**a**) or 100 (**b**) after infection of $p53^{+/+}$ or $p53^{-/-}$ mice with MHV68. **c,** Frequencies of infection and reactivation from panels **a,b. d-g,** Total cells in spleens from mock-infected or MHV68-infected $p53^{+/+}$ or $p53^{-/-}$ mice were quantified and analyzed by flow cytometry. B cells were gated as $CD19^{+}/B220^{+}$, GC B cells as $GL7^{+}/CD38^{lo}$ subset of B cells, and plasmablasts as $CD138^{+}/B220^{lo}$. Data represent means +/- SEM. No significant differences were present by Mann-Whitney unpaired *t* test.

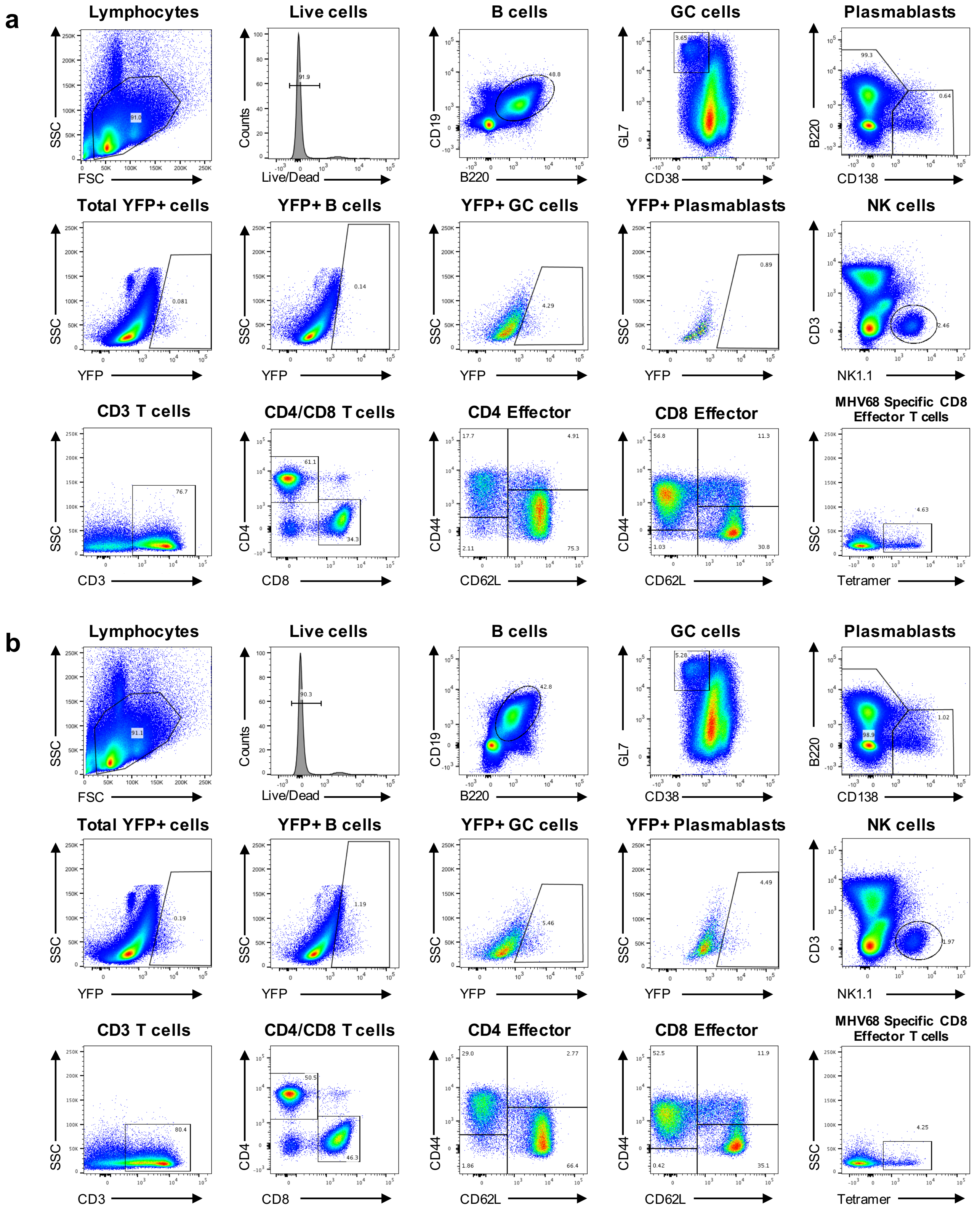


**Supplemental Figure 6: Representative flow cytometry plots demonstrating gating strategies for cellular phenotyping experiments. a,b,** p53$^{+/+}$ or p53$^{-/-}$ mice were infected intranasally with $10^4$ PFU of H2B-YFP MHV68 and sacrificed on day 16 post-infection. Splenocytes were analyzed by flow cytometry. Specific cell types were identified as follows: B cells (CD19$^+$/B220$^+$), GC B cells (GL7$^+$/CD38$^{lo}$) after gating on B cells, plasmablasts (CD138$^+$/B220$^{lo}$), T cells (CD3$^+$) followed by CD4$^+$ or CD8$^+$, effector T cells (CD62L$^-$/CD44$^+$) as CD4 or CD8 subgate, MHV68-specific effector T cells (ORF6-MHCI tet$^+$) as CD4 or CD8 effector subgate, and NK cells (CD3$^{lo}$/NK1.1$^+$). p53$^{+/+}$ mice are shown in **a**, and p53$^{-/-}$ mice are shown in **b**.

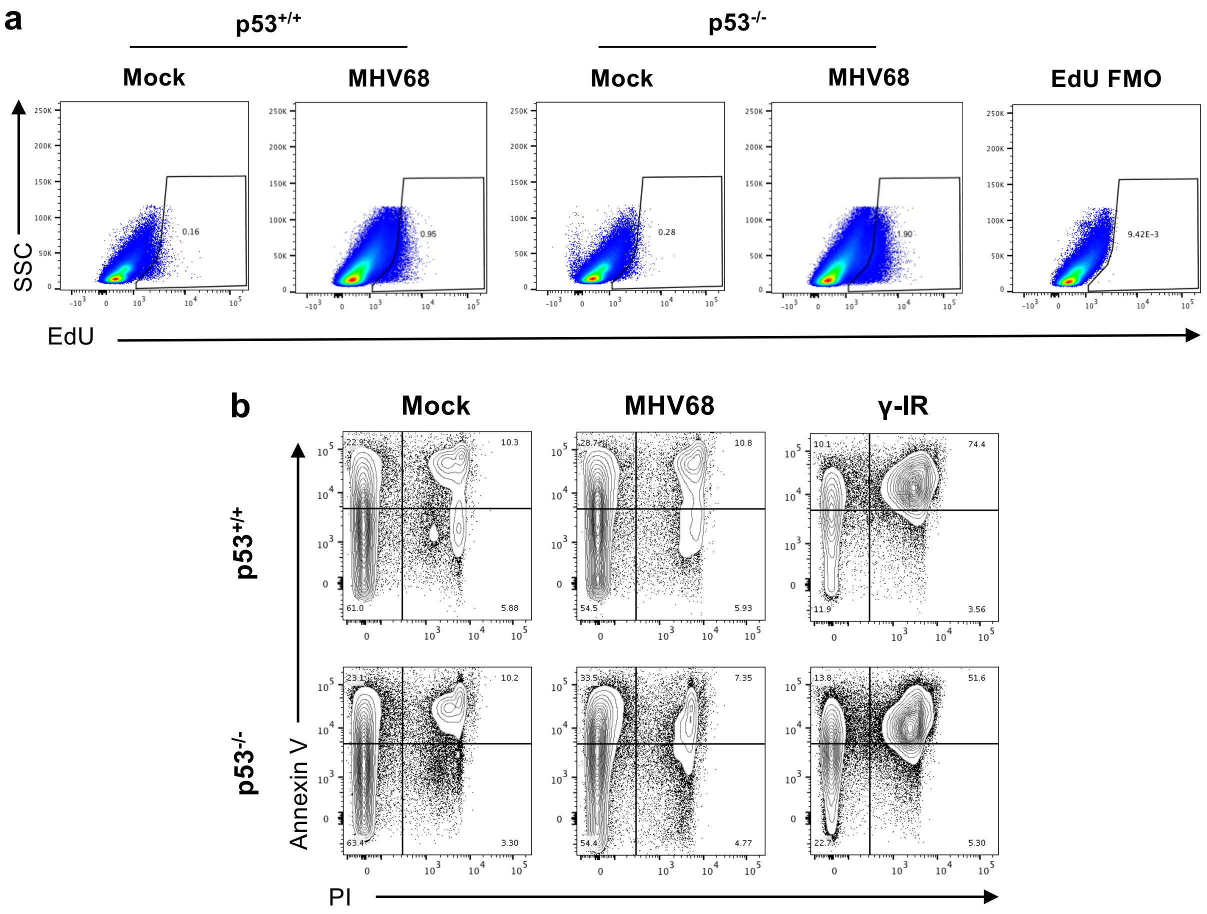


**Supplemental Figure 7: Representative flow cytometry plots demonstrating gating for EdU incorporation and Annexin V assays. a,b,** p53$^{+/+}$ or p53$^{-/-}$ mice were mock infected or infected intranasally with $10^4$ PFU of H2B-YFP MHV68 and sacrificed on day 16 post-infection. Splenocytes were analyzed by flow cytometry. Representative flow plots from experiments described in **Figure 4a** are shown in **a**. Representative flow plots from experiments described in **Figure 4b** are shown in **b**.

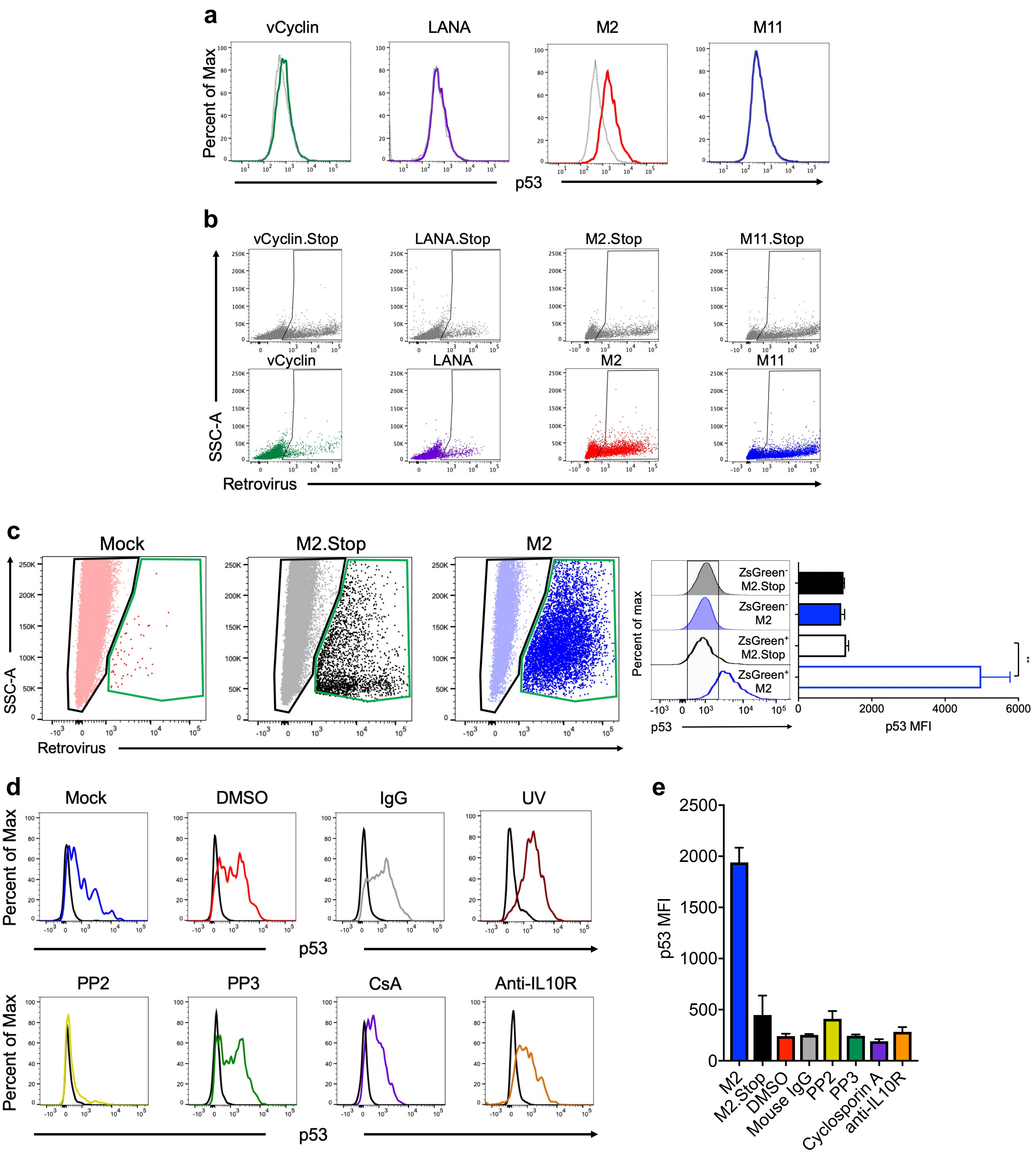


**Supplemental Figure 8: Representative flow cytometry plots demonstrating gating strategies and controls for p53 expression in B cells transduced with viral latency genes. a-e,** Representative flow cytometry plots and quantifications are shown for experiments described in **Figure 5**. Data in **a** show histograms in which p53 staining was overlaid for frame-shift stop control (dotted line) and viral latency gene (solid line) transduced B cells. Data in **b** show gating strategies to identify transduced cells. Data in **c** depict representative gating strategies for transduced and untransduced cells within single cultures and quantification of p53 staining for cells within these gates showing that p53 is induced specifically in M2-transduced B cells. of retrovirus-negative (black) and retrovirus-positive (green) populations. Data in **d** represent p53 staining in M2.Stop (black) or M2 (color) -transduced B cells in the presence of the indicated treatment. UV treatment serves as a positive control for p53 induction in non-transduced B cells. Mean fluorescence intensities of p53 staining were quantified for M2.Stop transduced B cells after the indicated treatments in **e**. M2-transduction is shown for comparison.

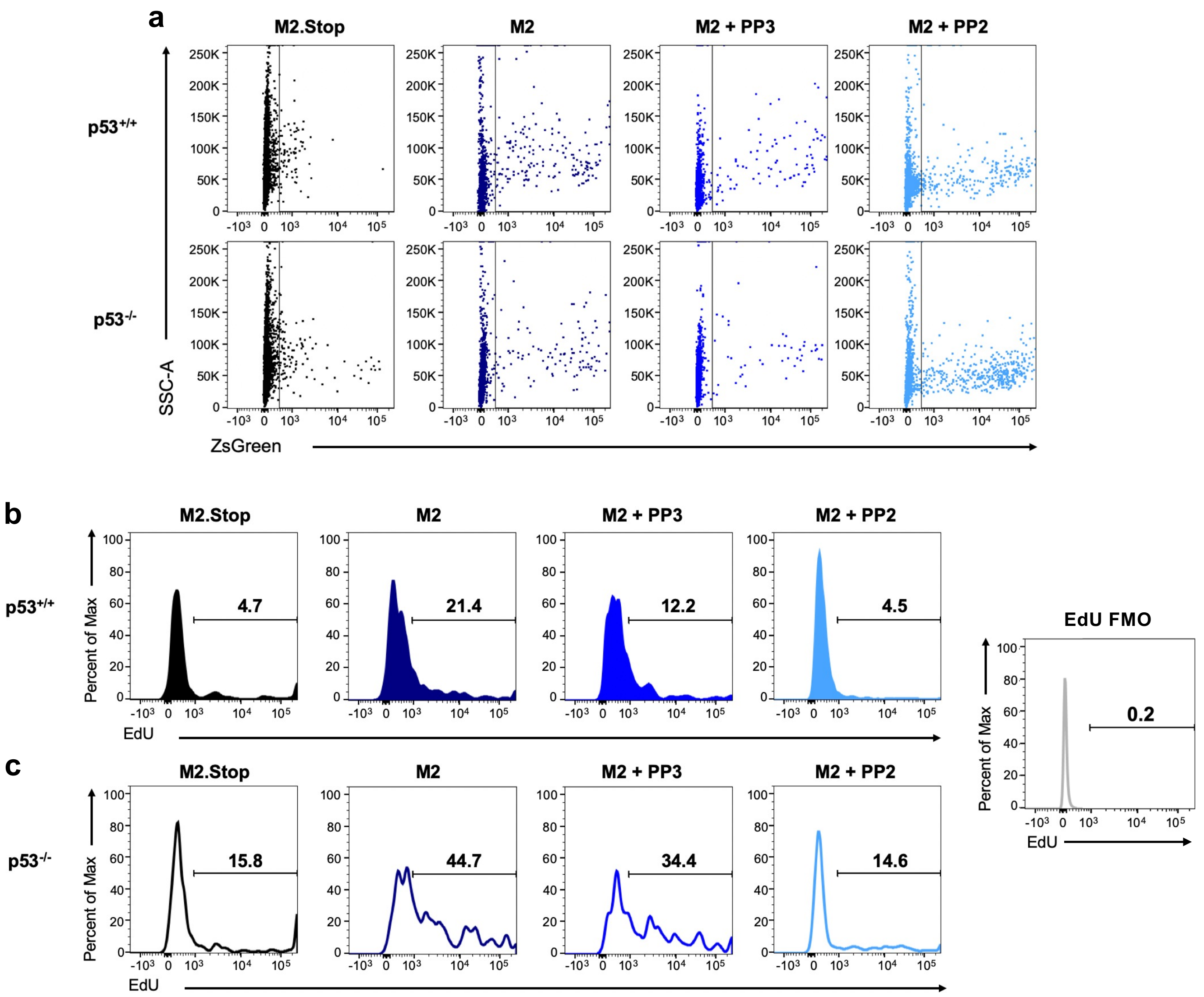


**Supplemental Figure 9: Representative flow cytometry plots showing EdU incorporation and retroviral transduction. a,** Flow cytometry plots showing ZsGreen detection 50 hours after transduction of $p53^{+/+}$ or $p53^{-/-}$ B cells. **b-c,** Transduced $p53^{+/+}$ or $p53^{-/-}$ B cells were labeled with EdU for 4 hours prior to harvest and EdU detection with Click chemistry. Representative flow cytometry plots from experiments described in **Figure 6** are shown in the presence or absence of the indicated treatments.

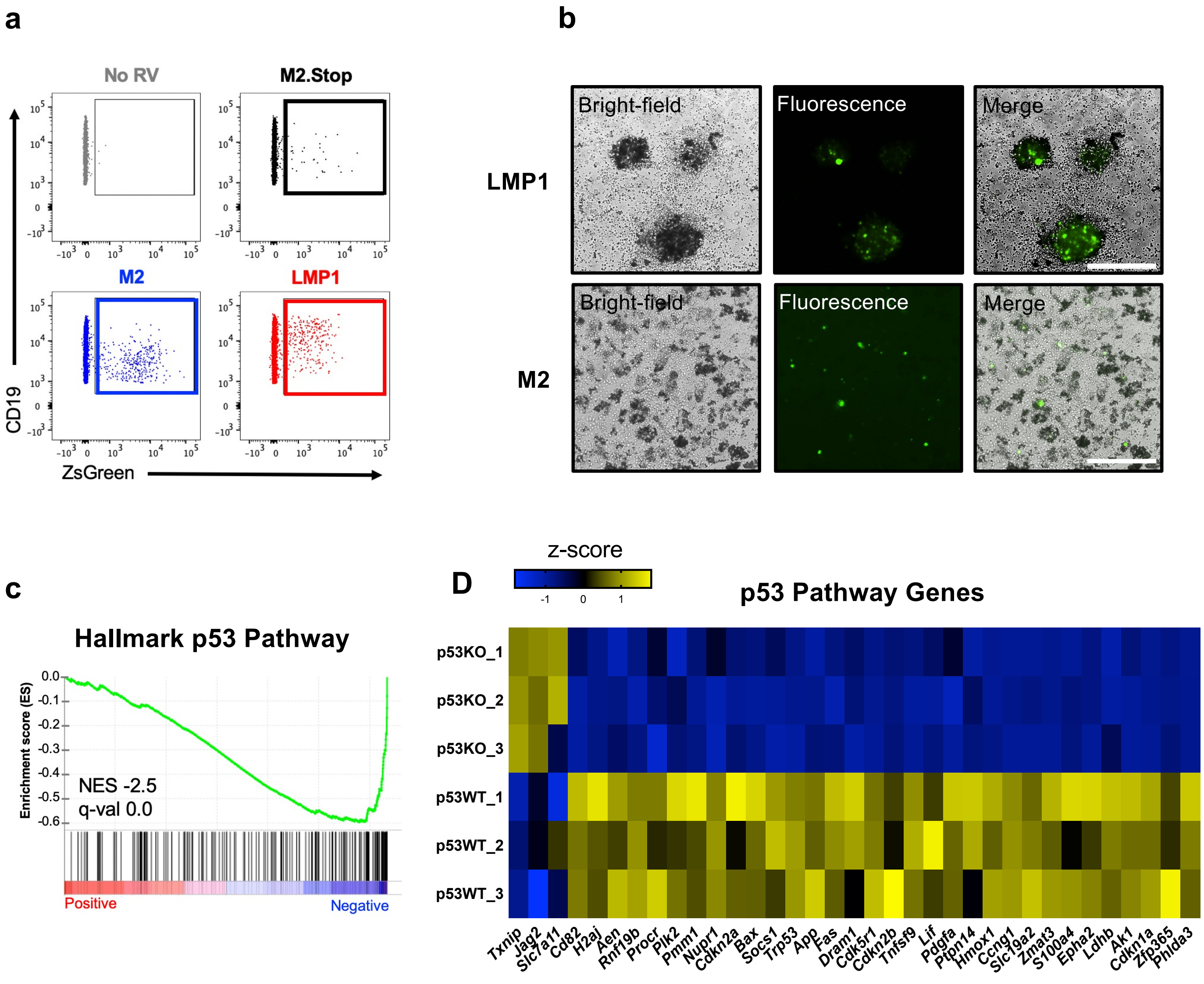


**Supplemental Figure 10: Representative flow cytometry plots for defining LMP1 effects on primary mouse B cells. a,** Representative flow cytometry plots showing Zs-Green detection following the indicated retroviral transductions corresponding to experiments described in **Figure 7**. **b,** fluorescence microscopy analysis showing surviving cell clusters 13 days after transduction of p53$^{-/-}$ B cells with M2 or LMP1-encoding retrovirus. Control transduction did not survive to this point. **c,** Gene set enrichment analysis (GSEA) and hallmark gene z-score heatmaps for the hallmark p53 gene set comparing RNA-seq data from p53+/+ and p53-/- B cells 4 days after transduction with LMP1 encoding retrovirus. The reference list was derived from Hallmark gene sets and compared with a pre-ranked list (by fold) of global average gene expression. Statistical scores are inset into the top right of analysis images. NES, normalized enrichment score. q-val, FDR-adjusted *p* value.

## M2

### $p53^{+/+}$

| Rank | Top Hallmark Gene Sets | NES |
|---|---|---|
| 1 | IL-2 STAT5 Signaling | 2.10 |
| 2 | Epithelial mesenchymal transition | 2.01 |
| 3 | Angiogenesis | 2.00 |
| 4 | TNFA Signaling via NFKB | 1.96 |
| 5 | Inflammatory Response | 1.94 |
| 6 | p53 Pathway | 1.91 |
| 1 | E2F Targets | -2.05 |
| 2 | G2M Checkpoint | -1.49 |

### $p53^{-/-}$

| Rank | Top Hallmark Gene Sets | NES |
|---|---|---|
| 1 | E2F Targets | 3.10 |
| 2 | G2M Checkpoints | 2.95 |
| 3 | MYC Targets | 1.97 |
| 1 | p53 Pathway | -2.55 |
| 2 | Epithelial mesenchymal transition | -2.26 |
| 3 | Angiogenesis | -2.05 |

### LMP1

### $p53^{+/+}$

| Rank | Top Hallmark Gene Sets | NES |
|---|---|---|
| 1 | MYC Targets V2 | 2.17 |
| 2 | MYC Targets V1 | 2.15 |
| 3 | p53 Pathway | 1.99 |
| 1 | Protein secretion | -1.16 |
| 2 | Notch Signaling | -0.99 |
| 3 | Bile Acid Metabolism | -0.95 |

### $p53^{-/-}$

| Rank | Top Hallmark Gene Sets | NES |
|---|---|---|
| 1 | Interferon Alpha Response | 2.12 |
| 2 | Interferon Gamma Response | 1.62 |
| 3 | KRAS Signaling | 1.40 |
| 1 | p53 Pathway | -2.51 |
| 2 | MYC Targets V2 | -2.39 |
| 3 | MYC Targets V1 | -2.35 |

**Supplemental Figure 11:** Top hallmark gene sets identified in GSEA for M2 and LMP1 expressing B cells. The tables show significantly up-regulated (yellow) or down-regulated (blue) hallmark gene sets for the indicated comparisons. The reference list was derived from Hallmark gene sets and compared with a pre-ranked list (by fold) of global average gene expression. NES, normalized enrichment score.
